# Dietary tryptophan enhances aryl hydrocarbon receptor activation and reduces colitis through microbial metabolism

**DOI:** 10.1101/2025.08.27.672750

**Authors:** Liam E Rondeau, Bruna B Da Luz, Dominic Haas, Pranshu Muppidi, Xuanyu Wang, Rebecca Dang, Gaston Rueda, Andrea Nardelli, Giada De Palma, Harry Sokol, Premysl Bercik, Alberto Caminero

**Affiliations:** Farncombe Family Digestive Health Research Institute, Department of Medicine, McMaster University, Hamilton, Canada; Université Paris-Saclay, INRAe, AgroParisTech, Micalis Institute, Jouy-en-Josas, France; Sorbonne Université, INSERM UMRS-938, Centre de Recherce Saint-Antoine, CRSA, AP-HP, Paris, France

**Keywords:** Tryptophan metabolism, aryl hydrocarbon receptor, inflammatory bowel disease, microbiota

## Abstract

**Background & Aim:** Disrupted microbial tryptophan metabolism and impaired aryl hydrocarbon receptor (AhR) activation are implicated in inflammatory bowel disease (IBD) pathogenesis. However, strategies to restore this pathway through diet or microbial modulation remain poorly defined. This study investigates how dietary tryptophan and human and mouse microbiota modulate metabolism, AhR activation, and intestinal inflammation in preclinical models.

**Methods:** Gnotobiotic mice colonized with microbiota of varying complexity or human fecal microbiota from ulcerative colitis (UC) patients and healthy controls were used to assess the impact of microbiota and dietary tryptophan supplementation on AhR activation and colitis severity. Chemically induced and spontaneous colitis models were investigated.

**Results:** IBD fecal samples showed reduced AhR activation compared to healthy controls, and fecal microbiota transplantation into germ-free mice demonstrated that impaired AhR is microbiota-dependent. Mice colonized with minimal microbiota had impaired microbial tryptophan metabolism, lower AhR activation, and worsened colitis severity compared to those colonized with complex microbiota. Dietary tryptophan supplementation in conventional and UC-humanized mice enhanced microbial production of AhR agonists, restored AhR activation, and reduced colitis severity in an AhR-dependent manner. Co-colonization with a tryptophan-metabolizing bacterium, *Clostridium sporogenes*, further improved tryptophan metabolism and colitis severity in mice with impaired microbial tryptophan metabolism.

**Conclusions:** Microbial tryptophan metabolism is critical for determining intestinal inflammation. Dietary tryptophan supplementation restores microbial metabolic pathways, mitigates colitis severity in preclinical models, and may address key metabolic deficiencies in IBD patients with impaired tryptophan metabolism. This study demonstrates the therapeutic potential of targeting microbial metabolism with diet in IBD management.

## INTRODUCTION

Inflammatory bowel disease (IBD), encompassing Crohn’s disease (CD) and ulcerative colitis (UC), is a chronic inflammatory disorder of the gastrointestinal tract with rising prevalence worldwide^1^. Beyond its debilitating symptoms, IBD imposes significant economic and societal burdens^2, 3^. While there have been advances in pharmacological therapies in recent years, treatment options remain expensive and limited, with many patients experiencing loss of efficacy or disease progression^4^. Dietary interventions represent a promising strategy to complement existing immunosuppressive treatments by addressing the factors underlying inflammation^5^.

The disruption of immune and epithelial homeostasis is a hallmark of IBD that is believed to be driven by interactions between gut microbiota and the mucosal immune system^6^. Alterations in gut microbiota composition and function^7^, as well as disrupted communication between host and microbiota have been demonstrated in IBD^6, 8–10^. The intestinal microbiota produces bioactive metabolites that regulate immune responses and barrier integrity, with those from dietary tryptophan having particular importance^11, 12^.

Tryptophan, an essential amino acid found in foods like poultry, is metabolized into compounds that activate the aryl hydrocarbon receptor (AhR), a transcription factor that regulates immune functions and gut homeostasis^11^. AhR promotes tissue repair and mucosal healing through the production of interleukin (IL)-22^13, 14^. While host enzymes metabolize the majority (90-95%) of tryptophan in the kynurenine pathway^15^, 4-6% of unabsorbed tryptophan undergoes microbial metabolism in the gut^16^, producing AhR agonists such as indole-3-acetic acid, tryptamine, and indole-3-propionic acid^17^. Bacterial genera involved in this process include *Lactobacillus*, *Clostridium*, and *Peptostreptococcus*^11, 18, 19^.

In IBD, reduced levels of AhR agonists and important microbial tryptophan metabolism genes have been observed in intestinal content of patients compared to healthy controls^12, 20, 21^, which is associated with decreased fecal AhR activation^12^. Expression of AhR is also downregulated in the intestinal tissue of IBD patients^22^, potentially contributing to impaired immune regulation and barrier dysfunction. These findings suggest a link between reduced AhR activity and IBD pathogenesis and raise questions about whether dietary or microbial interventions can restore AhR activation and reduce inflammation.

We investigated the interplay between dietary tryptophan, microbial metabolism, and AhR activation during intestinal inflammation. Using gnotobiotic models with microbiota of varying complexity, we demonstrated that impaired microbial tryptophan metabolism reduced AhR activation and worsened colitis severity. Enriching the diets of conventional mice with tryptophan enhanced microbial production of AhR agonists, leading to robust AhR activation and an AhR-dependent reduction in inflammation in multiple mouse models of colitis. Colonization with *Clostridium sporogenes*, a tryptophan-metabolizing bacterium, combined with dietary tryptophan supplementation, ameliorated colitis severity in mice with impaired tryptophan metabolism. Translational studies confirmed reduced tryptophan metabolite levels in IBD patients compared to healthy controls. Fecal microbiota transfer from humans to germ-free mice established the first direct evidence that AhR activation capacity and tryptophan metabolism profiles in human IBD are intrinsically linked to the gut microbiota. Importantly, dietary tryptophan supplementation restored AhR activation and alleviated intestinal inflammation in mice colonized with IBD-associated microbiota, highlighting the therapeutic potential of targeting microbial tryptophan metabolism with diet.

## RESULTS

### Intestinal microbiota mediates tryptophan metabolism and AhR activation

To investigate whether the capacity of different mouse microbiota to metabolize tryptophan influences AhR activation, germ-free (GF) mice were colonized with specific pathogen-free (SPF) or minimal microbiota (MM), a non-complex microbiota derived from Altered-Schaedler flora (Figure 1A). GF mice served as controls. Quantification of tryptophan metabolites in colon contents revealed that SPF mice had significantly higher levels of tryptophan-derived microbial metabolites, including tryptamine (TAM) and indole-3-propionic acid (I3PA) compared to MM-colonized mice and GF mice (Figure 1B). The total pool of microbial tryptophan metabolites was also substantially higher in SPF mice relative to MM mice (Figure 1C). As well, xanthurenic acid (XANA) and kynurenine (KYN), a host tryptophan metabolite produced by indoleamine 2,3-dioxygenase 1 (IDO1)^23,24^, were increased in colon contents of SPF mice (Figure 1B). GF mice had the lowest levels of all metabolites measured, confirming the essential role of gut microbiota in producing these AhR agonists.

**Figure 1.**
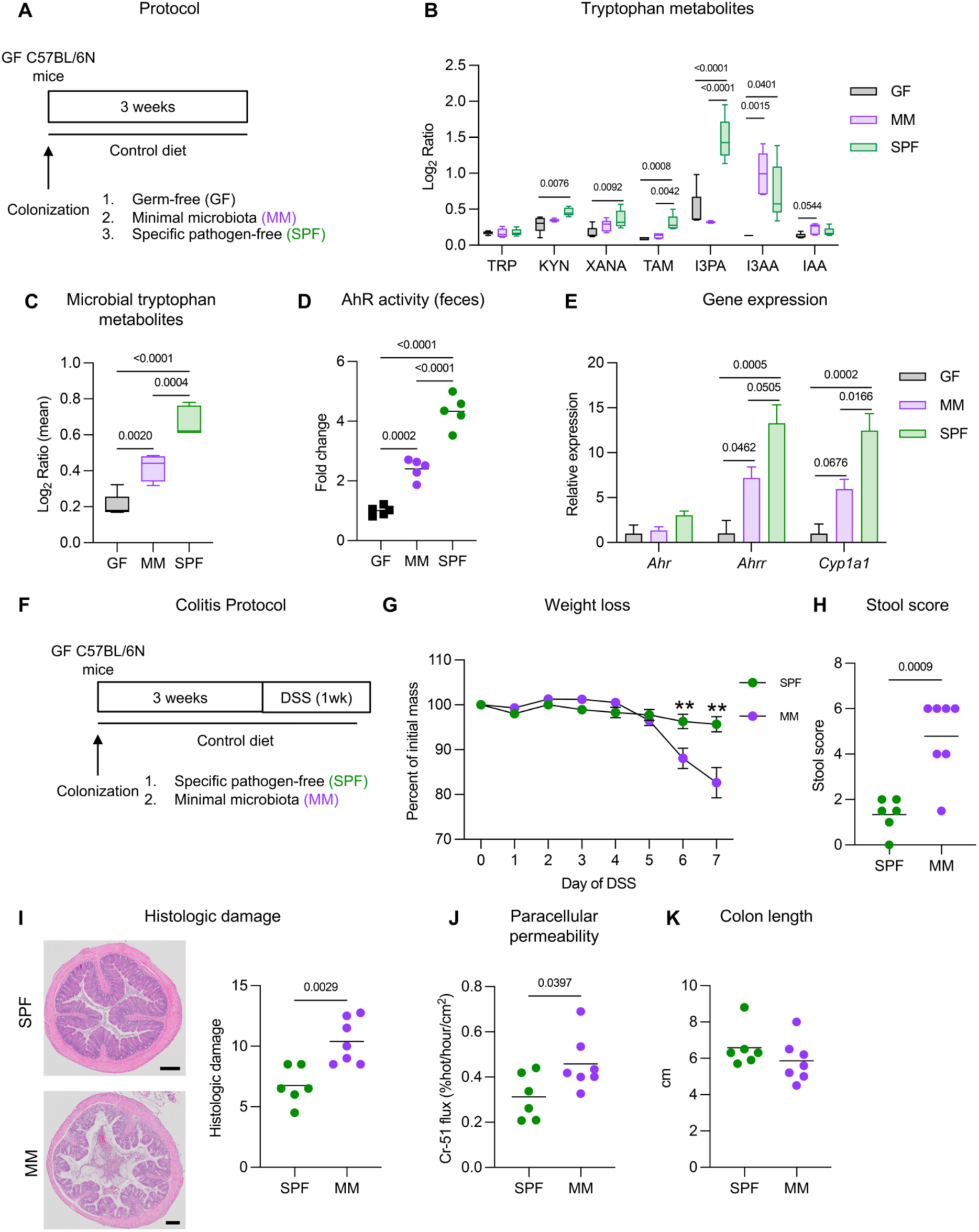
Microbiota mediates tryptophan metabolism, AhR activation, and colitis severity. **(a)** Protocol: Germ-free (GF) C57BL/6N mice were colonized with fecal/cecal slurries of minimal microbiota (MM), specific pathogen free (SPF) mice, or given a PBS vehicle control (GF) (n=5 mice per group). Tissues and intestinal contents were collected three weeks post-colonization. **(b)** Individual tryptophan metabolites in colon content comprising tryptophan (TRP), kynurenine (KYN), xanthurenic acid (XANA), tryptamine (TAM), indole-3-propionic acid (I3PA), indole-3-acetic acid (I3AA), and indoleacrylic acid (IAA). **(c)** Overall microbial tryptophan metabolites in colon content, including TAM, I3PA, I3AA, and IAA. **(d)** Fecal AhR activation capacity measured *in vitro*. **(e)** AhR pathway gene expression in colon tissue. **(f)** Colitis protocol: GF C57BL/6N mice were colonized with fecal/cecal slurries of minimal microbiota (MM) and specific pathogen-free (SPF) mice (n=6). Colitis was induced using 2.5% DSS for five days followed by a two-day washout. (n=6-7 mice per group). **(g)** Weight loss during colitis. **(h)** Composite stool score of stool blood and diarrhea at endpoint. **(i)** Histologic damage and representative distal colon sections from GF, MM, and SPF mice after colitis (H&E-stained; scale bars = 100 µm). **(j)** Paracellular permeability (Cr^51^-EDTA mucosal-to-serosal flux) of colon sections. **(k)** Colon length after colitis. Data are presented as median and interquartile range where whiskers extend to the min and max data points (B, C), as mean where each dot represents one mouse (D, H-K), and as mean + s.d. (G). Statistical analyses were performed using Kruskal-Wallis test with Dunn’s post-hoc test (B, C), one-way analysis of variance (ANOVA) with Tukey’s post-hoc test (D, E), and unpaired Student’s *t-*test (G-L). ** *p* < 0.01.

Consistent with these findings, fecal AhR activation capacity was highest in SPF-colonized mice, intermediate in MM-colonized mice, and lowest in GF controls, as measured by an *in vitro* AhR reporter assay (Figure 1D). Similarly, gene expression analysis of AhR pathway target genes in colon tissue showed that SPF mice had significantly elevated levels of *Ahr*, *Ahrr*, and *Cyp1a1* transcripts compared to MM mice, with GF mice exhibiting the lowest expression (Figure 1E). These results demonstrate that the complexity of the gut microbiota directly influences tryptophan metabolism and downstream AhR activation.

### Impaired tryptophan metabolism is associated with increased colitis susceptibility

To assess the functional consequences of altered microbiota-driven AhR activation in intestinal inflammation, we induced experimental colitis in MM- and SPF-colonized mice using dextran sulfate sodium (DSS) (Figure 1F). SPF-colonized mice exhibited reduced weight loss and lower stool scores (reduced severe diarrhea and stool blood), and less colon shortening compared to MM mice during colitis (Figure 1G–H). Histologic analysis revealed significantly greater mucosal damage in MM mice, with increased immune cell infiltration and epithelial disruption in distal colon sections (Figure 1I). Barrier function was also impaired in MM-colonized mice, as indicated by increased paracellular permeability (Cr-51 flux) measured *ex vivo* (Figure 1J).

To assess the molecular mechanisms involved with reduced AhR signaling and associated colitis severity, we analyzed colonic gene expression profiles after DSS treatment (Figure S1). MM-colonized mice exhibited significant upregulation of genes for pro-inflammatory cytokines and hall mark inflammatory mediators of colitis such as *Il1b*, *Tnf*, and *Myd88*. In contrast, SPF-colonized mice had increased expression of *Il22*, critical for promoting epithelial regeneration. Other cytokines regulated by AhR, including *Il21* and *Il17a* were also increased in SPF mice. *Il22ra2,* a receptor antagonist that inhibits the activity of IL-22^25^, was upregulated in MM mice compared to SPF mice during colitis, potentially limiting the protective effects of IL-22 and further contributing to the impaired barrier function and worsened colitis severity in MM mice. Therefore, the gut microbiota plays a critical role in regulating colonic immunity during inflammatory insults.

### Dietary tryptophan enrichment enhances AhR activation

To investigate the impact of dietary tryptophan enrichment on microbial tryptophan metabolism and AhR activation, SPF mice were fed either a control (CON) or high-tryptophan (HT) diet for three weeks (Figure 2A). Tryptophan supplementation increased tryptophan, and kynurenine (KYN), produced by host enzymes, in colon content (Figure 2B). Mice on the HT diet exhibited significantly elevated levels of TAM, I3PA, indole-3-acetic acid (I3AA), indoleacrylic acid (IAA), and overall microbial tryptophan metabolites compared to the CON-fed group (Figure 2B-C). These increases correlated with a 10-fold increase in fecal AhR activation (Figure 2D), and upregulated colonic expression of *Ahr, Ahrr, Cyp1a1, and Il22* (Figure 2E). The HT-fed mice also exhibited distinct microbiota profiles (Figure S3A-B), including increased abundance of Firmicutes members such as *Lactobacillus* and certain Clostridiales which have been associated with tryptophan metabolism, and reduced *Ruminococcaceae* and *Lachnospiraceae*. These results demonstrate that dietary tryptophan reaches the colon and can efficiently enhance microbial tryptophan metabolism, AhR activation, and downstream signaling in conventional SPF microbiota that has high basal tryptophan metabolic activity.

**Figure 2.**
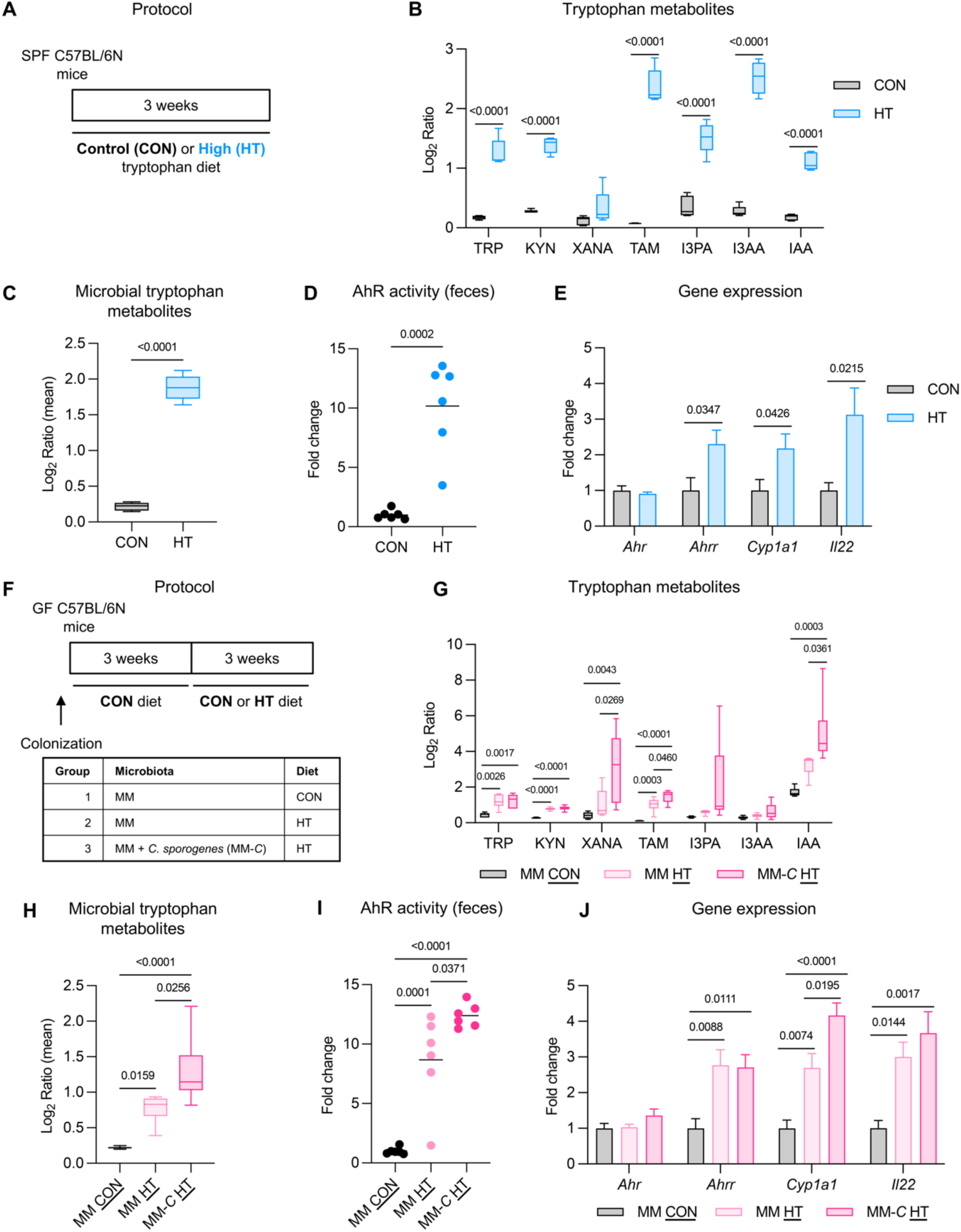
Dietary tryptophan enrichment activates AhR through efficient microbial tryptophan metabolism. **(a)** Protocol: Specific pathogen-free (SPF) C57BL/6N mice were fed a high tryptophan (HT) or control (CON) diet for three weeks. (n=6 mice per group) **(b)** Individual tryptophan metabolites in colon content including tryptophan (TRP), kynurenine (KYN), xanthurenic acid (XANA), tryptamine (TAM), indole-3-propionic acid (I3PA), indole-3-acetic acid (I3AA), and indoleacrylic acid (IAA). **(c)** Overall microbial tryptophan metabolites in colon content, including TAM, I3PA, I3AA, and IAA. **(d)** Fecal AhR activation capacity measured *in vitro*. **(e)** Gene expression of AhR pathway markers in colonic tissues. **(f)** Protocol: Germ-free (GF) C57BL/6 mice were colonized with minimal microbiota (MM) or MM supplemented with *Clostridium sporogenes* (ASF C). MM mice received either a control (CON) or high tryptophan (HT) diet, while MM-*C* mice received only the HT diet for three weeks. (n=6 mice per group). **(g)** Individual tryptophan metabolites in colon content comprising tryptophan (TRP), kynurenine (KYN), xanthurenic acid (XANA), tryptamine (TAM), indole-3-propionic acid (I3PA), indole-3-acetic acid (I3AA), and indoleacrylic acid (IAA) for MM and MM-*C*-colonized mice fed respective diets. **(h)** Overall microbial tryptophan metabolites in colon content, including TAM, I3PA, I3AA, IAA. **(i)** Fecal AhR activation capacity measured *in vitro*. **(j)** Gene expression of AhR pathway markers, hypothesized to be highest in MM-*C*-colonized mice fed the HT diet. Data are presented as median with interquartile range with whiskers extending to the min and max data points (B, C, G, H) mean (D, I), or mean ± s.d. (E, J). Statistical analyses were performed using Mann-Whitney U test (B, C), unpaired Student’s *t* test (D, E), one-way ANOVA with Tukey’s post-hoc test (J, K), and Kruskal-Wallis test with Dunn’s post-hoc test (H, I).

### Tryptophan metabolizing bacteria restore AhR activation in mice with impaired microbial tryptophan metabolism

To investigate whether a HT diet could restore AhR activation in microbiota with limited tryptophan metabolism capacity, GF mice were colonized with MM microbiota, which we demonstrated to have impaired tryptophan metabolism and reduced AhR activation. In addition, some mice were co-colonized with MM and *C. sporogenes* (MM-*C*), a bacterium isolated from healthy individuals and capable of efficiently metabolizing tryptophan into AhR agonists^26–29^, to assess whether *C. sporogenes* could compensate for the metabolic limitations of MM microbiota (Figure 2F). The capacity of *C. sporogenes* to activate AhR was demonstrated *in vitro*, and tryptophan metabolizing genes including indolelactate dehydrogenase (*fldH*), indolelactate dehydratase (*fldBC)*, acylCoA dehydrogenase (*AcdA*), and tryptophan decarboxylase (*TDC*) were also confirmed in our strain (Figure S2A-B). Fecal AhR activation was elevated in MM mice co-colonized with *C. sporogenes* before the HT diet was administered (Figure S2C).

MM-colonized mice fed the HT diet showed an increase in the levels of tryptophan, KYN, the microbial metabolite TAM (Figure 2G), as well as overall microbial tryptophan metabolites (Figure 2H) compared to those on the CON diet, indicating increased host and microbial tryptophan metabolic activity after tryptophan supplementation. MM are mostly colonized by *Ligilactobacillus* which may express tryptophan metabolism enzymes, albeit with significantly reduced affinity for tryptophan than other *Lactobacillus*^30^, which could potentially explain the slight increase in microbial tryptophan metabolites (Figure S2D). However, co-colonization with *C. sporogenes* and HT diet consumption resulted in significantly higher levels of KYN, TAM, and IAA compared to the CON diet, and XANA, TAM, and IAA compared to the HT diet alone (Figure 2G). Furthermore, there was a trend toward increased production of I3PA in MM-*C* mice on the HT diet. The *C. sporogenes* isolate used to colonize mice possesses genes encoding enzymes for IAA and I3PA production and is predicted to produce TAM, which may explain the enhanced levels of these microbial tryptophan metabolites in colon content (Figure S2). This enhancement was reflected in the overall microbial tryptophan metabolite pool, which was highest in MM-*C* mice on the HT diet (Figure 2H) and correlated with a significant increase in fecal AhR activation (Figure 2I) and upregulated colonic expression of *Ahrr*, *Cyp1a1,* and *Il22* compared to MM mice on the HT diet alone (Figure 2J).

These findings demonstrate that while dietary tryptophan alone can partially restore microbial tryptophan metabolism and AhR activation in MM-colonized mice, co-colonization with *C. sporogenes* provides additional benefit, further enhancing tryptophan metabolism and downstream signaling.

### Tryptophan dietary interventions alleviate colitis via AhR activation

To assess the therapeutic potential of dietary tryptophan in colitis, SPF mice were fed a CON or HT diet for three weeks, then colitis was induced with DSS, and the respective diets were maintained throughout colitis (Figure 3A). In SPF mice, feeding a HT diet for three weeks prior to DSS treatment significantly ameliorated disease severity compared to the CON diet. Mice fed HT exhibited lower histologic damage scores, with improved crypt architecture and reduced immune infiltration compared to CON-fed mice (Figure 3B). HT-fed mice also demonstrated lower stool scores (Figure 3C) and less colon shortening at endpoint (Figure 3D). The protective effects of HT were abrogated when an AhR antagonist (AhR⁻) was administered, indicating that these improvements were AhR-dependent. Weight loss during colitis was reduced in HT-fed mice compared to CON-fed mice, but this protective effect was not observed in the presence of the AhR antagonist (Figure 3E). HT diet feeding led to microbiota alterations and distinct clustering of colonic microbiota (Figure S3A-B), characterized by increased *Lactobacillus*, *Lactococcus*, and *Prevotellaceae*-UCG-001, and reduced *Lachnospiraceae, Tuzzerella, and Ruminococceae* (Figure S3C). The protection of an HT diet was also tested in hapten-induced and spontaneous models of colitis. In agreement with our findings in DSS colitis, HT diet reduced mucosal damage, the number of colonic ulcerations, and stool scores in mice exposed to TNBS, and reduced mucosal damage in IL-10-deficient mice (Figure S4).

**Figure 3.**
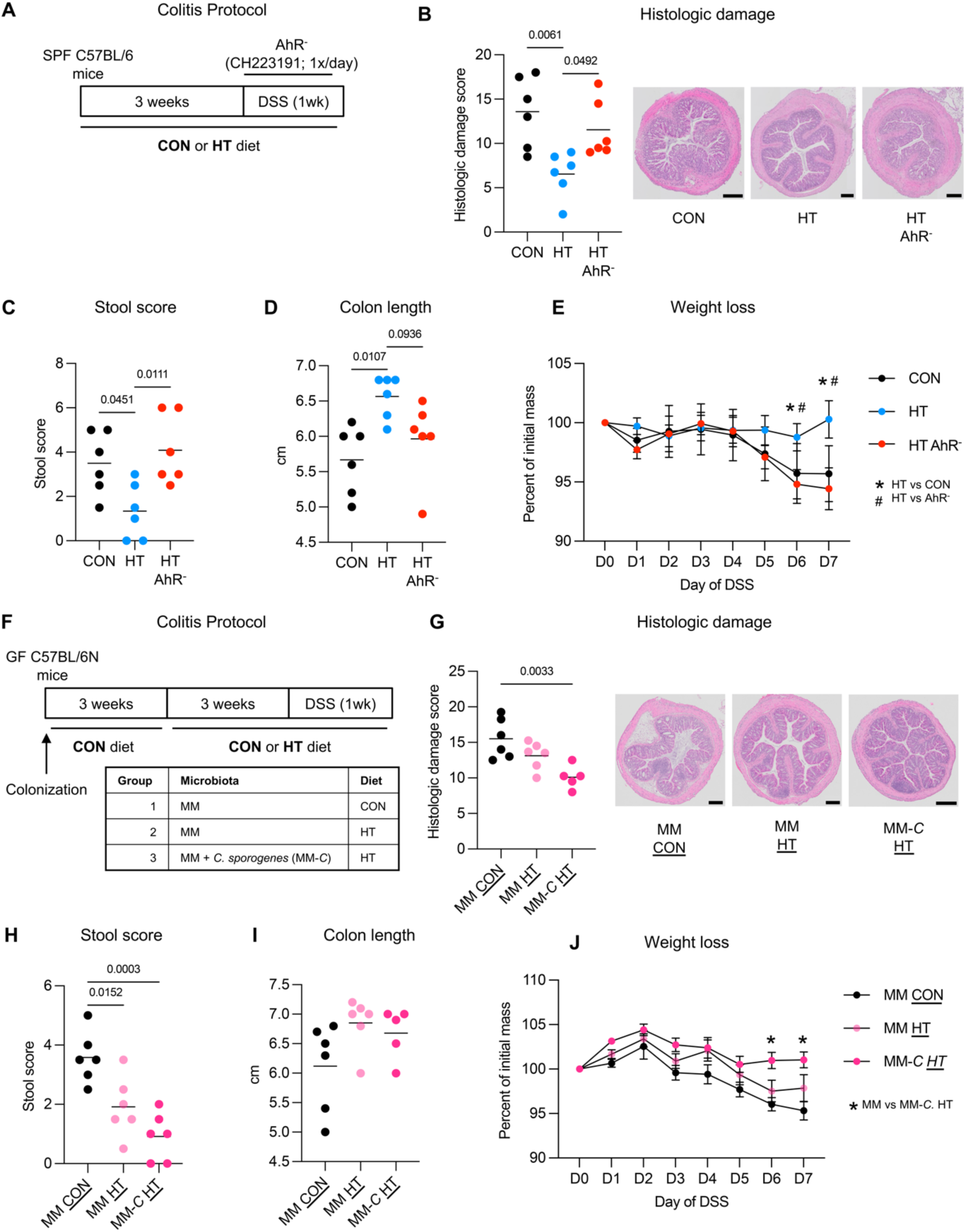
AhR activation by diet and microbes reduces colitis severity. **(a)** Colitis protocol: Specific pathogen-free (SPF) C57BL/6 mice were fed a high tryptophan (HT) or control (CON) diet for three weeks, then colitis was induced using 2.5% DSS for five days followed by a two-day washout. A subset of HT-fed mice received an AhR antagonist during colitis (AhR^-^; CH223191; 1×/day, oral gavage). (n=6 mice per group). **(b)** Histologic damage scores of CON, HT, and HT AhR⁻ mice, with representative distal colon sections (H&E-stained; scale bars = 200 μm). **(c)** Stool scores of CON, HT, and HT AhR⁻ mice, incorporating stool consistency and rectal bleeding. **(d)** Colon length of CON, HT, and HT AhR⁻ mice at endpoint. **(e)** Weight loss of CON, HT, and HT AhR⁻ mice during DSS treatment. **(f)** Colitis protocol: GF C57BL/6 mice were colonized with minimal microbiota (MM) or MM supplemented with *Clostridium sporogenes* (MM-*C*) and fed a GF conventional diet for three weeks. MM-colonized mice were fed either a control (CON) or high tryptophan (HT) diet, while MM-*C*-colonized mice were fed the HT diet only for three weeks before DSS treatment (2% DSS for five days followed by a two-day washout). (n=5-6 mice per group). **(g)** Histologic damage scores of mice, with representative distal colon sections (H&E-stained; scale bars = 200 μm). **(h)** Stool scores of MM CON, MM HT, and MM-*C* HT mice, incorporating stool consistency and rectal bleeding. **(i)** Colon length at endpoint. **(j)** Weight loss of mice during DSS treatment. Data are presented as mean (B-D, G-I) or mean ± s.d. (E, J). Statistical analyses were performed using one-way ANOVA with Tukey’s post-hoc test (B-E, G-J). *, # *p* < 0.05.

### Microbial and tryptophan interventions synergize to activate AhR and reduce inflammation in mice with impaired microbial tryptophan metabolism

Our results indicated that HT diet with or without *C. sporogenes* co-colonization increases AhR activation in MM-colonized mice. Thus, we next assessed the impact of increased tryptophan metabolism and AhR activity on DSS colitis severity in MM mice fed HT (MM HT) or co-colonized with *C. sporogenes* and fed HT (MM-*C* HT), compared to MM mice fed the CON diet (Figure 3F). In MM mice, which have impaired tryptophan metabolism capacity, dietary tryptophan alone (MM HT) moderately reduced colitis severity, but the combination of dietary tryptophan and *C. sporogenes* (MM-*C* HT) provided significantly greater protection. MM mice fed HT showed slight improvement in histologic damage compared to MM CON mice, but these effects were amplified when supplemented with *C. sporogenes* (MM-*C* HT), as indicated by reduced histologic damage scores and better-preserved tissue architecture (Figure 3G). While MM HT mice had reduced stool scores compared to CON-fed mice, MM-*C* HT mice exhibited the lowest stool scores among the groups (Figure 3H), alongside trends toward increased colon length (Figure 3I). Weight loss was attenuated in MM-*C* HT mice but not MM HT mice, compared to the MM CON groups during DSS treatment, further demonstrating the added benefit of microbial supplementation (Figure 3J).

Together, these findings underscore the critical role of dietary tryptophan and microbial tryptophan metabolism in promoting AhR activation and reducing colitis severity. Moreover, they highlight the potential of introducing tryptophan-metabolizing microbes to enhance tryptophan metabolism and AhR activation, particularly in contexts where microbial tryptophan metabolism is impaired.

### IBD patients have reduced AhR activation capacity and tryptophan metabolizing taxa

We next sought to confirm reduced fecal AhR activation capacity in IBD patients in a small Canadian cohort to support findings from previous studies^12^. Fecal samples from healthy controls (HC, n=10) and IBD patients (n=21) including CD (n=8) and UC, (n=13) were analyzed for their ability to activate AhR *in vitro*. Consistent with previous reports^12^, fecal AhR activation was significantly lower in IBD patients compared with HC, however it was not depleted suggesting the presence of tryptophan metabolizing taxa (Figure 4A).

**Figure 4.**
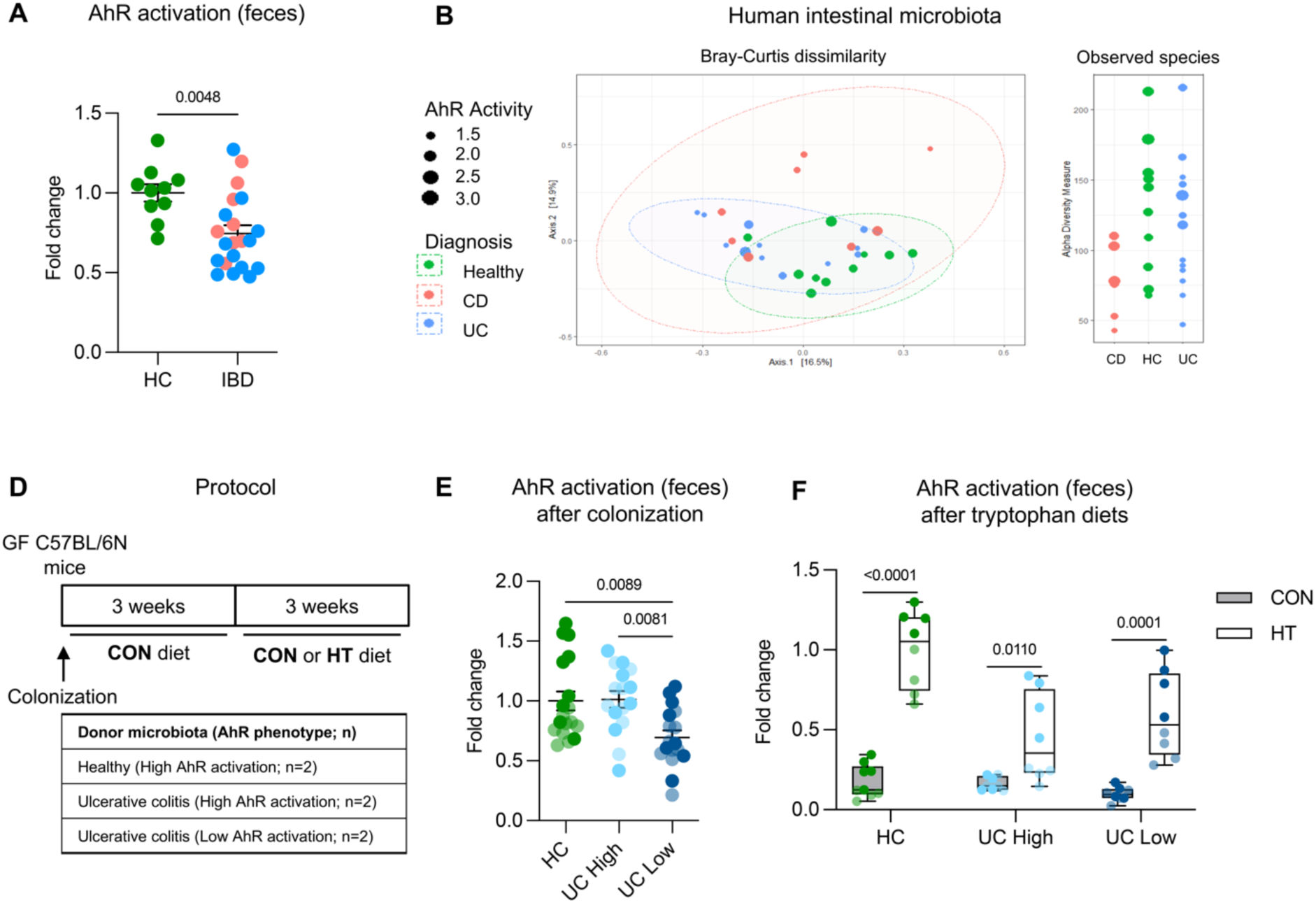
IBD microbiota impairs AhR activation via reduced microbial tryptophan metabolism, which is restored by a high-tryptophan diet in IBD humanized mice. **(a)** Fecal AhR activation in human fecal samples from healthy controls (HC; n=10) and IBD patients (n=21), including Crohn’s disease (CD; n=8; red dots) and ulcerative colitis (UC; n=13; blue dots). **(b)** Bray-Curtis dissimilarity and observed species richness of human fecal microbiota from HC, CD, and UC patients. Dot colors indicate donor diagnosis (HC, CD, or UC), and dot size reflects the level of fecal AhR activation in the corresponding sample. **(c)** Relative abundance of bacterial taxa implicated in tryptophan metabolism in HC and IBD patients. **(d)** Mouse colonization and tryptophan intervention protocol: Germ-free (GF) C57BL/6 mice were colonized with fecal microbiota from HC donors (high AhR activation, n=2), UC patients with high AhR activation (UC High, n=2), or UC patients with low AhR activation (UC Low, n=2). Colonized mice were maintained on either a control (CON) or high tryptophan (HT) diet for three weeks after initial colonization (n=8-9 mice per group). **(e)** Fecal AhR activation in GF mice after colonization with HC, UC High, or UC Low microbiota. Dot shading represents individual human donors within each group. **(f)** Fecal AhR activation in GF mice after colonization and subsequent consumption of CON or HT diet. Data are presented as mean ± s.e.m. where each dot represents one human or mouse (A, E) or median with interquartile range with whiskers extending to min and max values (F). Statistical analyses were performed using Student’s *t* test (A) and one-way ANOVA with Tukey’s post-hoc test (E, F).

Fecal microbiota between HC and IBD patients presented some differences. Bray-Curtis dissimilarity demonstrated slight clustering of samples by diagnosis, while observed species richness tended to be reduced in CD compared to HC and UC (Figure 4B). Importantly, IBD samples with higher AhR activation tended to exhibit microbiota profiles more similar to HC, suggesting a potential link between microbial composition and AhR activation capacity (Figure 4B). We also examined the relative abundance of bacterial taxa associated with tryptophan metabolism and observed a reduction in *Clostridiales,* an order associated with many tryptophan metabolizing taxa^11, 31^, in IBD patients compared to HC (Figure S5). Altogether, our findings confirm that the microbiota of IBD patients is distinct in composition and functional capacity, with fewer tryptophan metabolizing bacteria and diminished AhR activation.

### Impaired AhR activation in mice colonized with IBD microbiota is restored by a high tryptophan diet

We next investigated whether fecal microbiota from HC or IBD patients with varying AhR activation capacities could transfer AhR phenotypes to GF mice and whether a HT diet could enhance AhR activation in these mice. GF mice were colonized with fecal microbiota from HC (n=2 donors) or UC patients, categorized into high AhR activation (UC High, n=2 donors) and low AhR activation (UC Low, n=2 donors) groups, then maintained on either a CON or HT diet for three weeks (Figure 4D).

Following colonization (Figure S6), fecal samples from mice colonized with HC microbiota exhibited significantly higher AhR activation compared to those colonized with microbiota from UC low patients, with a similar level observed in mice colonized with HC and UC high microbiota (Figure 4E). These results demonstrate that the capacity for AhR activation is transferable through the microbiota, providing strong evidence that reduced AhR activation observed in IBD patients is microbiota dependent.

To evaluate the HT diet’s potential to restore AhR activation in mice with IBD-associated impaired microbial tryptophan metabolism, we analyzed fecal AhR activation in colonized mice after three weeks of dietary intervention with CON or HT diets. In all donor groups, the HT diet significantly increased AhR activation compared to the CON diet. Importantly, the HT diet was particularly effective in restoring AhR activation in mice colonized with UC low microbiota, increasing activation levels by nearly fourfold compared to the CON diet (Figure 4F). Altogether, while impaired AhR activation is mediated by IBD microbiota, dietary tryptophan supplementation can enhance the production of AhR agonists, even in hosts with low baseline tryptophan metabolism capacity.

### Tryptophan supplementation restores microbial tryptophan metabolism and alleviates colitis severity in IBD-humanized mice

To investigate the therapeutic potential of dietary tryptophan supplementation in restoring AhR activation and reducing colitis severity in IBD microbiota colonized mice, GF C57BL/6N mice were colonized with UC Low microbiota (characterized by low AhR activation capacity). After three weeks of stabilization on a GF diet, mice were divided into two groups and fed either a CON or HT diet for an additional three weeks. Colitis was then induced during diet consumption with DSS (Figure 5A).

**Figure 5.**
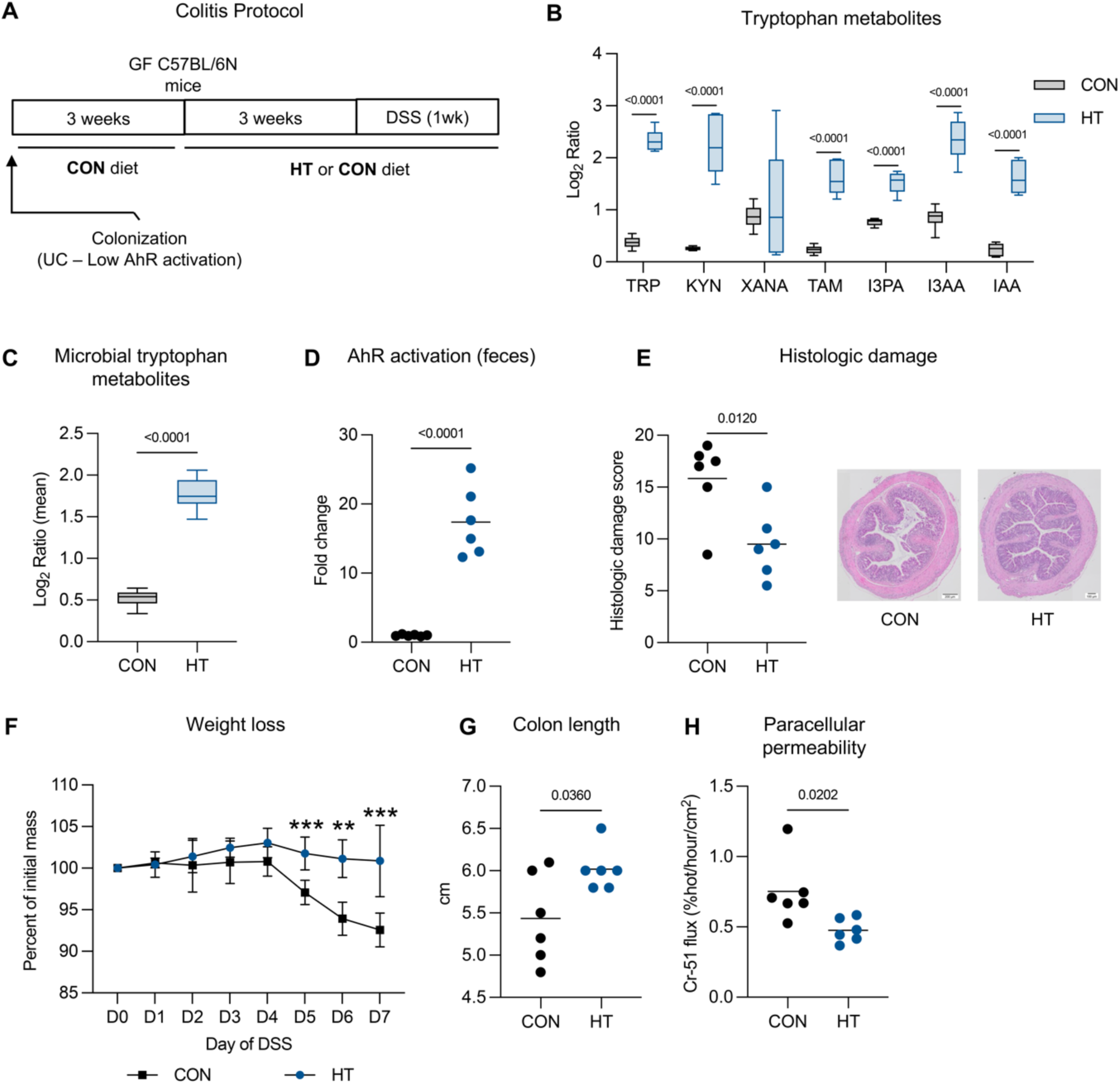
Tryptophan supplementation activates AhR and reduces inflammation in humanized IBD mice. **(a)** Colitis protocol: Germ-free (GF) C57BL/6 mice were colonized with microbiota of ulcerative colitis (UC) patient with low AhR activation, then fed a GF diet for three weeks. Mice were then fed to a control (CON) or high tryptophan (HT) diet for three weeks, then colitis was induced using 2.5% DSS for five days followed by a two-day washout. **(b)** Individual tryptophan metabolites in colon content, including tryptophan (TRP), kynurenine (KYN), xanthurenic acid (XANA), tryptamine (TAM), indole-3-propionic acid (I3PA), indole-3-acetic acid (I3AA), and indoleacrylic acid (IAA). **(c)** Overall microbial tryptophan metabolites in colon content, including TAM, I3PA, I3AA, and IAA. **(d)** Fecal AhR activation measured in vitro. **(e)** Histologic damage scores of colon tissue with representative images (H&E-stained; scale bars = 200 μm). **(f)** Weight loss during DSS treatment. **(g)** Colon length at endpoint. **(h)** Paracellular permeability of colonic tissue measured as Cr-51 flux (%/hour/cm²). Data are presented as median with interquartile range where whiskers extend to the min and max data points (B, C) or mean ± s.e.m. (D-H). Statistical analyses were performed using Mann-Whitney U tests for metabolomics (B, C) and unpaired Student’s t-tests for all other panels (D-H). ** *p* < 0.01, *** *p* < 0.001.

Metabolomic analysis revealed that the HT diet significantly increased levels of host tryptophan metabolites and several microbial tryptophan metabolites, including KYN, TAM, I3AA, IAA, and indoxyl (IND) (Figure 5B). Additionally, overall microbial tryptophan metabolites were elevated in HT-fed mice (Figure 5C), indicating enhanced microbial tryptophan metabolism. This increase in microbial metabolites was accompanied by a significant boost in fecal AhR activation capacity in HT-fed mice compared to CON-fed mice (Figure 5D).

The HT diet also ameliorated colitis severity, as evidenced by significantly reduced histologic damage scores (Figure 5E) and weight loss during DSS treatment (Figure 5F). HT-fed mice experienced less colon shortening during colitis compared to CON-fed mice, indicative of reduced inflammation and tissue damage (Figure 5G). Barrier function was also improved in HT-fed mice, as shown by reduced paracellular permeability (Figure 5H).

Altogether, these results confirm that dietary tryptophan can enhance microbial tryptophan metabolism and reduce colitis severity in IBD-humanized mice, even in the context of microbiota with impaired AhR activation capacity. The increase in microbial tryptophan metabolites suggest that dietary tryptophan increases both the metabolic activity of the existing microbiota and the population of tryptophan metabolizing taxa.

## DISCUSSION

Dietary interventions represent a promising strategy for managing IBD by targeting gut microbiota metabolism and host immunity. Tryptophan, an essential amino acid with an estimated average requirement of 4 mg/kg/day (280 mg/day for a 70-kg adult)^32^, is metabolized by host and microbial enzymes into AhR agonists that help maintain intestinal homeostasis^33^. Here, we show that microbial tryptophan metabolism is linked to AhR activation, and supplementation of dietary tryptophan can restore microbial metabolism and mitigate colitis severity. Our work suggests that dietary recommendations involving tryptophan consumption should be considered for IBD patients.

The role of dysregulated tryptophan metabolism and AhR activation in IBD has been previously reported^12, 21, 22^. Our work confirms and extends previous clinical observations that IBD patients exhibit lower fecal AhR activation and microbial tryptophan metabolites than HC^12, 21, 22^. Importantly, transplant of fecal microbiota from IBD patients to GF mice recapitulated this defect, with UC-associated microbiota with low tryptophan metabolism capacity driving a significant reduction in intestinal AhR activation compared to HC microbiota. These results establish a direct link between human IBD-associated microbiota and reduced AhR activation, resolving questions about a microbial contribution to this phenotype. We observed reduced levels of *Clostridiales*, an order enriched in tryptophan-metabolizing taxa^27^, in IBD patients, supporting the rationale for restoring microbial tryptophan metabolism through dietary and microbial interventions.

The importance of the intestinal microbiota in tryptophan metabolism and homeostasis has been established^33^. We confirmed that the complexity of gut microbiota plays an important role in tryptophan metabolism and immune regulation. Mice colonized with MM had reduced colonic microbial tryptophan metabolites, impaired AhR activation, and worsened colitis severity compared to mice conventional SPF colonized mice. This aligns with earlier work showing that *Card9*-deficient mice—genetically predisposed to colitis— harbor microbiota with defective tryptophan metabolism, leading to reduced AhR activation and impaired IL-22 production^12^. Interestingly, we found that microbiota composition not only influences microbial production of AhR agonists, including TAM and I3PA, but also host-derived metabolites including KYN and XANA—a protective host metabolite whose reduced levels predict relapse in IBD remission^24^—which were higher in mice colonized with complex SPF microbiota. These data show the importance of microbiota for determining tryptophan metabolism and immune responses during intestinal inflammation.

Leveraging the microbiota’s metabolic capacity for therapeutic purposes through dietary supplementation is a key objective in the field. Others have shown that *in vitro* incubation of fecal samples with tryptophan leads to increased production of AhR agonists like I3PA and indole-3-lactic acid in a tryptophan-dose-dependent manner, demonstrating the importance of the interplay between diet and gut microbes in producing tryptophan metabolites^29, 34^. We found that dietary tryptophan supplementation enhanced microbial metabolism of tryptophan in conventional mice, increasing metabolites including I3AA, TAM, and I3PA, while boosting AhR activation and reducing colitis severity. Dietary tryptophan also promoted the expansion of tryptophan metabolizing taxa, like *Lactobacillus*, suggesting that elevated substrate availability may enrich beneficial microbes^17, 35, 36^. Notably, even mice colonized with UC-associated microbiota exhibiting low AhR activation capacity responded to dietary intervention alone, with sufficient microbial tryptophan metabolism capacity to use the substrate, activate AhR, and reduce inflammation. However, in MM mice with limited microbial tryptophan metabolism mostly attributed to *Ligilactobacillus*, dietary tryptophan alone provided only partial protection. Co-colonization with *C. sporogenes*, previously shown to metabolize tryptophan into I3PA and other important AhR agonists in a tryptophan-dose-dependent manner^19, 27, 29, 31^, amplified dietary tryptophan’s effects, restoring AhR activation to achieve improvement in inflammation. Tryptophan supplementation also increased host production of XANA in MM mice after co-colonization with *C. sporogenes*, a potential benefit beyond the contribution of microbial metabolism as it may reduce colitis severity through AhR^24^. Although our use of an AhR antagonist confirms the role of this receptor in mediating the protective effects of dietary tryptophan, the involvement of other receptors such as the pregnane X receptor, involved in regulating intestinal barrier function through some microbial tryptophan metabolites^37^, cannot be ruled out entirely.

Tryptophan supplementation has been shown to be safe in humans and effective at boosting microbial indole production and AhR activation in healthy individuals^38^, supporting its potential as a therapeutic strategy for IBD patients. Indeed, the method of administration and the complexity of the dietary matrix are determinants for tryptophan reaching the colon for microbial metabolism^39^. While our findings demonstrate the efficacy of dietary interventions on their own in hosts with sufficient microbial metabolic capacity, they also suggest that combined strategies using targeted microbial supplementation may be beneficial in cases where there is severely impaired microbial function. Indeed, understanding the interactions between gut microbes and different dietary components, such as fibre or fat, and their effect on the balance between microbial and host tryptophan metabolism should also guide the optimization of dietary interventions as others have shown that excess kynurenine and serotonin production may be deleterious^23, 29, 40, 41^.

The protective effects of AhR activation on barrier integrity and inflammation make it a strong target for therapeutic interventions. Here, we establish the impact of microbial metabolism of dietary tryptophan and subsequent AhR activation in mitigating inflammation during colitis. By demonstrating that tryptophan and microbial supplementation increase microbial tryptophan metabolism and the production of host-derived metabolites like XANA, even in microbiota with impaired tryptophan metabolic capacity, we suggest its potential as a therapeutic strategy for IBD. Importantly, we provide the first direct evidence that the microbiota is responsible for AhR activation in the human gut, with fecal microbiota transplantation experiments confirming that IBD microbiota drives impaired AhR activation. Our work also reinforces the potential for targeted microbial supplementation, such as with *C. sporogenes* or other tryptophan metabolizing bacteria like *Lactobacillus*^17^, to restore tryptophan metabolism in combination with diet. Clinical studies that evaluate combinatorial diet and microbial approaches are warranted to improve IBD management.

## MATERIALS AND METHODS

### Human study design and patient population

This study involved secondary laboratory analysis of previously collected human fecal samples and associated de-identified metadata. No new recruitment or clinical data collection was performed. Participants were originally recruited through a collaboration with Dr. Premysl Bercik at McMaster University. Fecal samples were collected from individuals with a confirmed diagnosis of IBD and from healthy controls (HC). IBD diagnosis was established based on a combination of clinical presentation, endoscopic findings and histopathological evaluation in accordance with accepted diagnostic guidelines. Fecal samples were collected during follow-up visits or prior to surveillance colonoscopy and stored under anaerobic conditions before being frozen at −80°C until analysis. Inclusion criteria for IBD patients were a confirmed diagnosis of CD or UC with age ≥18 years. Healthy controls were individuals undergoing colonoscopy for colorectal cancer screening, with no evidence of organic disease, and no history of IBD. Exclusion criteria for all participants included recent gastrointestinal infection and use of antibiotics or probiotics within four weeks of sample collection. Fecal samples were collected from 22 individuals with IBD (8 with CD and 13 with UC) and 10 HC. Demographic and clinical data, including age, sex, weight, medication use, and medical history, were recorded for all participants and are shown in Table S1. All participants provided written informed consent before enrollment, and the original study was approved by the Hamilton Integrated Research Ethics Board (HIREB # 15311). The study was registered on ClinicalTrials.gov (ID: NCT06532110; https://clinicaltrials.gov/study/NCT06532110). AhR activation was measured in fecal samples to assess its relationship with microbial tryptophan metabolism. Microbiota analysis was performed for HC and IBD patient groups. Samples were used for colonization of germ-free (GF) mice.

### Mouse study designs

Six-to-eight-week-old specific pathogen-free (SPF) and eight-to-twelve-week-old GF C57BL/6N mice were used to investigate the roles of microbiota, dietary tryptophan, and AhR activation in colitis. GF mice were colonized with distinct microbial communities or human fecal microbiota to evaluate microbial tryptophan metabolism and its downstream effects on AhR activation and inflammation. Colonization was performed via oral gavage with microbial suspensions prepared from: (i) SPF microbiota, (ii) minimal microbiota (MM), a low complexity well-defined microbial community of eight species (Altered Schaedler flora-derived microbiota) with low tryptophan metabolism, (iii) MM co-colonized with *Clostridium sporogenes*, or (iv) human fecal microbiota obtained from healthy controls (HC), ulcerative colitis patients with high AhR activation (UC High), or low AhR activation (UC Low). Colonized mice were housed under GF conditions and fed a control GF diet for three weeks to allow for microbiota stabilization.

For dietary experiments, mice were transitioned to either a control (CON) or high-tryptophan (HT) diet for three weeks, after which they either underwent colitis (and maintained on the experimental diets), or were terminated. In subsets of mice, colitis was induced using dextran sulfate sodium (DSS) and 2,4-6 trinitrobenzene sulfonic acid (TNBS) to model intestinal inflammation and investigate the effects of AhR activation and microbial tryptophan metabolism. Additionally, SPF IL-10-deficient BALB/C mice were fed a CON or HT diet for 12 weeks before intestinal inflammation was assessed.

Each individual mouse was considered the experimental unit for all analyses and the total number of mice used per group and total sample sizes are provided in figure captions. No animals were excluded based on pre-established criteria. All mice that completed the study were included in the final analysis. No animals or data points were excluded from analysis unless indicated in figure legends. Primary outcome measures included colon histology scores and AhR activation levels, which guided sample size decisions. Throughout, sample sizes were calculated based on prior experiments using gnotobiotic mouse models of colitis^42, 43^.

### Mice

Specific-pathogen-free (SPF) C57BL/6N mice of both sexes were purchased from Taconic and housed at McMaster University’s Central Animal Facility (CAF) under SPF conditions. For colonization and gnotobiotic experiments, GF C57BL/6N mice were generated through two-stage embryo transfer and bred under GF conditions in the Axenic Gnotobiotic Unit (AGU) at the Farncombe Family Digestive Health Research Institute. MM microbiota was derived from the cecal and colonic contents of MM C3H-HeN mice maintained in the AGU for more than 15 generations, and originally purchased as mice harbouring Altered Schaedler flora (eight bacterial strain community) from Taconic^42^. Six-to-eight-week-old IL-10-deficient BALB/C mice were purchased from Taconic and housed under SPF conditions. All mice were provided unlimited access to food and water and maintained on a 12-hour light/12-hour dark cycle. All experiments were approved by the McMaster University Animal Care Committee and conducted in accordance with the Animal Utilization Protocol #22-26. Ethical guidelines for animal research were strictly followed, and mice were matched by sex and age. Mice were randomly assigned to treatment groups without a formal randomization sequence. Fecal sample collection was performed in a consistent order and at similar times of day. Outcome assessment was blinded to group allocation; blinding during diet administration was not feasible due to housing logistics. All mice were allowed to acclimate for 1 week before experimental procedures unless colonized GF.

### Gnotobiotic mouse colonization

GF mice were maintained in sterile isolators to ensure GF conditions prior to colonization. Microbial suspensions were prepared by diluting fecal and cecal content 1:10 in sterile phosphate-buffered saline (PBS) under anaerobic conditions. Mice were colonized via oral gavage with 200 μL of the prepared slurry. Colonization groups included: (i) MM, (ii) SPF microbiota prepared from recently euthanized donor mice, or (iii) PBS as a vehicle control. MM-colonized mice were further co-colonized with *C. sporogenes* (10⁹ CFU/mouse), grown anaerobically in brain heart infusion (BHI) medium. Human fecal microbiota was also used to colonize GF mice, using fecal material from HC, UC High, or UC Low donors. After colonization, mice were housed under GF conditions and fed a control GF diet for three weeks to allow microbiota stabilization.

### Tryptophan diet formulations

Mice were provided *ad libitum* access to iso-caloric custom-formulated versions of the Envigo Teklad AIN-93M diet (TD.00102), modified to adjust tryptophan content. The control diet (CON) contained no additional tryptophan (0.14% total tryptophan by weight), while the high-tryptophan diet (HT) included 0.8% supplemental tryptophan by weight (0.94% total tryptophan by weight). The detailed composition of each diet is presented in Table S2.

### AhR antagonist (CH223191)

To assess the role of AhR signaling in response to diet and microbial intervention during inflammation, select groups of mice were treated with the AhR antagonist CH223191. Mice received CH223191 (10 mg/kg body weight) by oral gavage once daily during the five-day DSS colitis induction phase. The antagonist was prepared fresh daily by dissolving CH223191 in dimethyl sulfoxide (DMSO) and diluting with sterile phosphate-buffered saline (PBS) to achieve a final DMSO concentration of 0.1%. Mice in control groups received vehicle-only gavages.

### Metabolomic analysis of mouse intestinal content

Colon contents were collected from mice at the endpoint of experiments and flash-frozen on dry ice prior to storage at −80°C. For metabolomic profiling, samples were processed by The Metabolomics Innovation Centre (TMIC) at the University of Alberta. Metabolites were extracted by homogenizing approximately 50 mg of colon content in 500 µL of LC-MS-grade methanol:water (4:1, v/v) using a bead beater at 5 m/s for 15 seconds. The homogenates were incubated at −20°C for 10 minutes, centrifuged at 15,000 g for 10 minutes, and the supernatants were collected, dried under vacuum, and reconstituted in 30 µL of LC-MS-grade water.

Chemical isotope labeling was performed to enhance the detection and quantitation of metabolites. Samples were labeled with 12C2 reagents for individual samples and 13C2 reagents for pooled reference samples, following the manufacturer’s protocol. The labeled samples were analyzed using high-performance liquid chromatography coupled with tandem mass spectrometry (LC-MS/MS) on a Thermo Scientific Vanquish LC system coupled to a Bruker Impact II QTOF mass spectrometer. Chromatographic separation was achieved on a C18 column using a gradient of 0.1% formic acid in water and acetonitrile.

Untargeted metabolomics focused on quantifying tryptophan and its microbial and host metabolites, including kynurenine, tryptamine, indole-3-propionic acid, indole-3-acetic acid, indoleacrylic acid, and xanthurenic acid. Metabolites were identified and quantified using TMIC’s NovaMT Metabolite Database. Data normalization to pooled references ensured cross-sample comparability, and metabolite quantities were presented as peak intensity ratios (Log2 scale).

Metabolomic data processing was performed using IsoMS Pro software for peak detection, quantification, and quality control. Overall microbial metabolites were calculated by averaging the Log_2_ ratios for all microbial metabolites of each sample. The data are presented as median with interquartile range, with whiskers extending to the minimum and maximum values of Log2 ratios or weighted summation, and inter-group differences in tryptophan metabolism were assessed using the Mann-Whitney U test or Kruskal-Wallis test.

### Measurement of AhR activity

AhR activation of mouse and human fecal samples was measured using an AhR-expressing luciferase reporter assay as previously described^12, 17, 44^. Briefly, H1L1.1c2 cells, which contain the dioxin response element-driven firefly luciferase reporter plasmid pGudLuc1.1, were seeded in 96-well plates and stimulated with stool filtrates in triplicate for 24 hours. Luciferase activity was measured after incubation using a luminometer. All values were normalized based on the cytotoxicity of the samples using the Lactate Dehydrogenase Activity Assay (Promega). All data are expressed as fold change of the control group.

To measure AhR activation by bacteria, individual *C. sporogenes* colonies were picked and cultured anaerobically in either BHI medium or tryptophan medium (Sigma). Cultures were incubated overnight at 37°C, and media controls were included to account for background effects. After incubation, bacterial cultures were centrifuged at 12,000 g for 10 minutes to collect the supernatants, which were filtered through 0.22 μm filters to remove residual cells. Filtered supernatants were applied to H1L1.1c2 cells and AhR activation was quantified as described above.

### Gene expression

Total RNA was extracted from RNAlater-stabilized (Invitrogen) mouse colon tissue samples using the RNeasy Mini Kit (Qiagen) following the manufacturer’s protocol. Global expression of inflammatory genes was assessed using the NanoString nCounter platform with the Mouse Inflammation Panel (254 genes). For genes not included in the NanoString panel, quantitative RT-PCR (RT-qPCR) was conducted using iScript Reverse Transcriptase (Bio-Rad) and SsoFast EvaGreen Supermix (Bio-Rad) with specific mouse primers (Table S3). Gene expression analysis was performed using the 2^−ΔΔCt^ quantification method for RT-qPCR data. Key genes in the AhR signaling pathway, including *Ahr*, *Ahrr*, *Cyp1a1*, and *Il22* were amplified and analyzed and normalized to *Gapdh*.

### Identification of tryptophan metabolizing genes

DNA was extracted from a bacterial pellet by boiling individual colonies and sequenced on the Illumina MiSeq platform (Illumina, San Diego, CA). We then used IDBA-hybrid using the NCBI accession ASM102020v1 as the reference genome of *C. sporogenes* to obtain the contigs against which we used NCBI blastn to search for similarities to our tryptophan metabolism enzyme sequences of interest.

### Induction and assessment of DSS colitis

Intestinal injury was induced in mice by administering 2% or 2.5% dextran sulfate sodium (DSS; 36,000–50,000 MW; MP Biomedicals LLC) in drinking water for five days, followed by two days on normal drinking water before sacrifice. Mice that did not receive DSS were given water. DSS intake was monitored daily throughout all experiments.

Colitis severity was evaluated by measuring colon length, body-weight loss, stool consistency, and stool blood. Stool consistency and blood were each scored on a scale of 0–3, as previously described^45^, then summed for a stool score.

### Induction and assessment of TNBS colitis

Trinitrobenzenesulfonic acid (TNBS) colitis was induced in 12-week-old SPF C57BL/6N mice as previously described. Briefly, mice were anesthetized with isoflurane (4%) and administered TNBS (2% in 30% ethanol) rectally using a polyethylene catheter inserted into the colon. Mice were monitored for body weight loss and mortality over 72 hours. At the endpoint, colons were excised, rinsed with saline, and evaluated for macroscopic damage using a scoring system which assigns scores based on severity and extent of inflammation and tissue damage. Score: 0- No damage. 1- Hyperemia and thickening of bowel wall without ulcers. 2- Hyperemia and thickening of bowel wall without ulcers. 3- One site of ulceration without bowel wall thickening. 4- Two or more sites of ulceration or inflammation. 5- 0.5 cm of inflammation and major damage. 6 to 10- 1 cm of major damage. The score is increased by one for every 0.5 cm of damage observed, up to a maximum of 10. Colon length and stool score were measured and scored as described in the DSS procedures above.

### IL-10-deficient colitis

Six-to-eight-week-old IL-10-deficient mice that develop spontaneous colitis were fed either CON or HT diets upon arrival for 12 weeks. At endpoint, histologic damage of the distal colon was assessed as a measure of inflammation.

### Evaluation of microscopic damage after colitis

For histologic analysis of DSS, TNBS, and IL-10-deficient colitis, tissue samples were collected from the distal colon, placed in 10% formalin, and embedded in paraffin. All tissues were then stained with filtered hematoxylin (Sigma-Aldrich) and 1% eosin Y solution (Sigma-Aldrich) and visualized for histological evaluation of morphology under light microscope (Olympus, ON, Canada). Two sections of the distal colon were evaluated for evidence of inflammation by two blinded observers. Microscopic damage on a scale of 0–21 was determined, based on a modified scoring system^46^. The following parameters were assessed: a) Presence of polymorphonuclear cells and erythrocytes in luminal exudate (absent, score = 0, scant = 1, moderate = 2, dense = 3); b) Epithelial damage and desquamation (no pathological change, score = 0, mild regenerative change = 1, moderate with patchy desquamation = 2, severe with diffuse desquamation = 3); c) Acute inflammatory infiltrate of the mucosa (PMNs) (absent, score = 0, scant = 1, moderate = 2, dense = 3); d) Mononuclear cell infiltrate of the mucosa (absent, score = 0, one small aggregate = 1, more than one aggregate = 2, several aggregates = 3); e) Goblet cell depletion (abundant number of goblet cells, score = 0, mild/moderate = 1, severe = 2); Cryptitis and crypt abscesses (absent, score = 0, isolated cryptitis = 1, diffuse cryptitis = 2, crypt abscesses = 3); Architectural damage (no pathological change, score = 0, mild and isolated architectural change = 1, moderate change = 2, severe and diffuse changes = 3); Edema (absent, score = 0, present = 1). The sum of all the parameters was used to determine the histologic damage score.

### Intestinal permeability

Colonic paracellular permeability (^51^Cr-EDTA mucosal-to-serosal flux) was evaluated *ex vivo* using proximal colon sections and the Ussing chamber technique, as previously described^47^. Sections of proximal colon (1.5cm) were collected, opened along the mesenteric border, and mounted into an Ussing chamber. The chamber exposed 0.25 cm^2^ of tissue surface area to 4 ml of circulating oxygenated Krebs buffer containing 10 mM glucose (serosal side) and 10 mM mannitol (mucosal side) maintained at 37°C aerated with 95% O_2_ and 5% CO_2_. The flux rate of ^51^Cr-EDTA from mucosa to serosa was quantified in samples using a liquid scintillation counter and expressed as % hot sample/h/cm^2^. The Ussing chamber system was from Physiologic Instruments (Reno, NV, USA).

### Microbiota analysis

Mouse fecal pellets and human feces were collected. DNA was extracted from samples and the hypervariable V3 regions of the 16S rRNA gene were amplified with polymerase chain reaction (PCR) using Taq polymerase (Life Technologies, Carlsbad, CA), as previously described^46^. Primers targeting the V3 region (v3f_ R1-5’-ATTACCGCGGCTGCTGG; v3r_ R2-5’-CCTACGGGNGGCWGCAG) were used. Reverse primers included six-base pair barcodes to allow multiplexing samples. Purified PCR products were sequenced using the Illumina MiSeq platform by the McMaster Genomics facility. Primers were trimmed from the obtained sequences with Cutadapt software^48^, and processed with Divisive Amplicon Denoising Algorithm 2 (DADA2; version 1.14.0) using the trained SILVA reference database (version 138.1)^49^. A phylogenetic tree of the sequences was calculated using FastTree 2^50^, and data was explored using the phyloseq package (version 1.30.0) in R (version 3.6.2)^51^. After data cleanup for human fecal samples, a total of 1,593,097 reads were obtained with a minimum of 6,108 and maximum of 87,578 with an average of 52,455 reads per sample. For mouse samples, a total of 14, 367,637 reads were obtained, with a minimum of 7660 and maximum of 212,725, and an average of 118,247 reads per sample. Alpha-diversity was measured using observed species. Beta-diversity was calculated on normalized data using the Bray-Curtis dissimilarity method. The originated matrix was ordinated using non-metric multidimensional scaling (NMDS).

### Statistical analysis

All variables were analyzed with SPSS version 18.0 (SPSS Inc., USA) and GraphPad Prism 9 (GraphPad Software, USA). Categorical variables are presented as numbers with percentages, while quantitative variables are expressed as means ± standard deviation (s.d.) for parametric data, and as medians with interquartile ranges for nonparametric data. Parametric data are depicted as dot plots with each dot representing an individual mouse or biological replicate, or as bar charts. Normal distribution was determined by D’Agostino–Pearson omnibus normality test, Shapiro–Wilk test, and Kolmogorov– Smirnov test with Dallal–Wilkinson–Lillie correction. The one-way analysis of variance (ANOVA) test was used to evaluate differences between more than two groups with a parametric distribution and Tukey’s post-hoc correction was applied. Student’s *t*-test (two-tailed) was performed to evaluate the differences between two independent groups as appropriate. Data with non-normal distribution were evaluated with Kruskal–Wallis test with Dunn’s post-hoc test for more than two groups, and Mann–Whitney test for two independent groups. Wald test was used for statistical analysis of gene expression by Nanostring nSolver 2.5. A *P* value < 0.05 was selected to reject the null hypothesis by two-tailed tests. Information regarding specific *P* values, value of *n*, and how data are presented can be found in figure legends.

## Author contributions

Conceptualization, LER, AC; Methodology, LER, BBDL, AC; Investigation, LER, BBDL, DH, PM, XYW, DR, GDP; Writing–Original Draft Preparation, LER, AC; Writing–Review and Editing, LER, BBDL, GDP, HS, PB, AC; Supervision, AC; Funding acquisition, AC, PB.

## Abbreviations

AhR: aryl hydrocarbon receptor
CD: Crohn’s disease
CON: control diet
GF: germ-free
HC: healthy control
HT: high tryptophan diet
IBD: inflammatory bowel disease
IAA: indoleacrylic acid
I3AA: indole-3-acetic acid
I3PA: indole-3-propionic acid
IL-10: interleukin-10
IND: indoxyl
KYN: kynurenine
MM: minimal microbiota
MM-C: minimal microbiota co-colonized with Clostridium sporogenes
SPF: specific pathogen-free
TAM: tryptamine
TRP: tryptophan
UC: ulcerative colitis
XANA: xanthurenic acid

## Synopsis

This study demonstrates that dietary tryptophan restores microbial production of AhR agonists and reduces colitis severity by activating the homeostatic AhR pathway, showing a therapeutic strategy to reduce intestinal inflammation through diet and microbial metabolism.

## Acknowledgements

The authors thank the staff of the Axenic Gnotobiotic Unit at McMaster University, Joe Notorangelo, Michael Rosati, and Sarah Armstrong for their assistance with mouse care. We also thank the McMaster Genomics Facility, Laura Rosati, and Michelle Shah for technical support with 16S rRNA gene sequencing and NanoString assays. We thank M.S. Denison for providing the H1L1.1c2 cell line and The Metabolomics Innovation Centre and Dr. Liang Li for metabolomic analyses.

## REFERENCES

1. Ng SC, Shi HY, Hamidi N, et al. Worldwide incidence and prevalence of inflammatory bowel disease in the 21st century: a systematic review of population-based studies. Lancet 2017;390:2769–2778.

2. Bernstein CN, Longobardi T, Finlayson G, et al. Direct medical cost of managing IBD patients: a Canadian population-based study. Inflamm Bowel Dis 2012;18:1498–508.

3. Kuenzig ME, Lee L, El-Matary W, et al. The Impact of Inflammatory Bowel Disease in Canada 2018: Indirect Costs of IBD Care. J Can Assoc Gastroenterol 2019;2:S34–S41.

4. Curro D, Pugliese D, Armuzzi A. Frontiers in Drug Research and Development for Inflammatory Bowel Disease. Front Pharmacol 2017;8:400.

5. Fitzpatrick JA, Melton SL, Yao CK, et al. Dietary management of adults with IBD - the emerging role of dietary therapy. Nat Rev Gastroenterol Hepatol 2022;19:652–669.

6. de Souza HS, Fiocchi C. Immunopathogenesis of IBD: current state of the art. Nat Rev Gastroenterol Hepatol 2016;13:13–27.

7. Lloyd-Price J, Arze C, Ananthakrishnan AN, et al. Multi-omics of the gut microbial ecosystem in inflammatory bowel diseases. Nature 2019;569:655–662.

8. Zheng D, Liwinski T, Elinav E. Interaction between microbiota and immunity in health and disease. Cell Res 2020;30:492–506.

9. Lee M, Chang EB. Inflammatory Bowel Diseases (IBD) and the Microbiome-Searching the Crime Scene for Clues. Gastroenterology 2021;160:524–537.

10. Lu Y, Li X, Liu S, et al. Toll-like Receptors and Inflammatory Bowel Disease. Front Immunol 2018;9:72.

11. Lamas B, Natividad JM, Sokol H. Aryl hydrocarbon receptor and intestinal immunity. Mucosal Immunol 2018;11:1024–1038.

12. Lamas B, Richard ML, Leducq V, et al. CARD9 impacts colitis by altering gut microbiota metabolism of tryptophan into aryl hydrocarbon receptor ligands. Nat Med 2016;22:598–605.

13. Qiu J, Heller JJ, Guo X, et al. The aryl hydrocarbon receptor regulates gut immunity through modulation of innate lymphoid cells. Immunity 2012;36:92–104.

14. Veldhoen M, Hirota K, Westendorf AM, et al. The aryl hydrocarbon receptor links TH17-cell-mediated autoimmunity to environmental toxins. Nature 2008;453:106–9.

15. Cervenka I, Agudelo LZ, Ruas JL. Kynurenines: Tryptophan’s metabolites in exercise, inflammation, and mental health. Science 2017;357.

16. Wyatt M, Greathouse KL. Targeting Dietary and Microbial Tryptophan-Indole Metabolism as Therapeutic Approaches to Colon Cancer. Nutrients 2021;13.

17. Lamas B, Hernandez-Galan L, Galipeau HJ, et al. Aryl hydrocarbon receptor ligand production by the gut microbiota is decreased in celiac disease leading to intestinal inflammation. Sci Transl Med 2020;12.

18. Wlodarska M, Luo C, Kolde R, et al. Indoleacrylic Acid Produced by Commensal Peptostreptococcus Species Suppresses Inflammation. Cell Host Microbe 2017;22:25–37 e6.

19. Wikoff WR, Anfora AT, Liu J, et al. Metabolomics analysis reveals large effects of gut microflora on mammalian blood metabolites. Proc Natl Acad Sci U S A 2009;106:3698–703.

20. Morgan XC, Tickle TL, Sokol H, et al. Dysfunction of the intestinal microbiome in inflammatory bowel disease and treatment. Genome Biol 2012;13:R79.

21. Franzosa EA, Sirota-Madi A, Avila-Pacheco J, et al. Gut microbiome structure and metabolic activity in inflammatory bowel disease. Nat Microbiol 2019;4:293–305.

22. Monteleone I, Rizzo A, Sarra M, et al. Aryl hydrocarbon receptor-induced signals up-regulate IL-22 production and inhibit inflammation in the gastrointestinal tract. Gastroenterology 2011;141:237–48, 248 e1.

23. Chajadine M, Laurans L, Radecke T, et al. Harnessing intestinal tryptophan catabolism to relieve atherosclerosis in mice. Nat Commun 2024;15:6390.

24. Michaudel C, Danne C, Agus A, et al. Rewiring the altered tryptophan metabolism as a novel therapeutic strategy in inflammatory bowel diseases. Gut 2023;72:1296–1307.

25. Zenewicz LA. IL-22 Binding Protein (IL-22BP) in the Regulation of IL-22 Biology. Front Immunol 2021;12:766586.

26. Caminero A, Herran AR, Nistal E, et al. Diversity of the cultivable human gut microbiome involved in gluten metabolism: isolation of microorganisms with potential interest for coeliac disease. FEMS Microbiol Ecol 2014;88:309–19.

27. Dodd D, Spitzer MH, Van Treuren W, et al. A gut bacterial pathway metabolizes aromatic amino acids into nine circulating metabolites. Nature 2017;551:648–652.

28. Stickland LH. Studies in the metabolism of the strict anaerobes (genus Clostridium): The chemical reactions by which Cl. sporogenes obtains its energy. Biochem J 1934;28:1746–59.

29. Sinha AK, Laursen MF, Brinck JE, et al. Dietary fibre directs microbial tryptophan metabolism via metabolic interactions in the gut microbiota. Nat Microbiol 2024;9:1964–1978.

30. Montgomery TL, Eckstrom K, Lile KH, et al. Lactobacillus reuteri tryptophan metabolism promotes host susceptibility to CNS autoimmunity. Microbiome 2022;10:198.

31. Williams BB, Van Benschoten AH, Cimermancic P, et al. Discovery and characterization of gut microbiota decarboxylases that can produce the neurotransmitter tryptamine. Cell Host Microbe 2014;16:495–503.

32. Lieberman HR, Agarwal S, Fulgoni VL, 3rd. Tryptophan Intake in the US Adult Population Is Not Related to Liver or Kidney Function but Is Associated with Depression and Sleep Outcomes. J Nutr 2016;146:2609S–2615S.

33. Roager HM, Licht TR. Microbial tryptophan catabolites in health and disease. Nat Commun 2018;9:3294.

34. Koper JE, Troise AD, Loonen LM, et al. Tryptophan Supplementation Increases the Production of Microbial-Derived AhR Agonists in an In Vitro Simulator of Intestinal Microbial Ecosystem. J Agric Food Chem 2022;70:3958–3968.

35. Bender MJ, McPherson AC, Phelps CM, et al. Dietary tryptophan metabolite released by intratumoral Lactobacillus reuteri facilitates immune checkpoint inhibitor treatment. Cell 2023;186:1846–1862 e26.

36. Cervantes-Barragan L, Chai JN, Tianero MD, et al. Lactobacillus reuteri induces gut intraepithelial CD4(+)CD8alphaalpha(+) T cells. Science 2017;357:806–810.

37. Flannigan KL, Nieves KM, Szczepanski HE, et al. The Pregnane X Receptor and Indole-3-Propionic Acid Shape the Intestinal Mesenchyme to Restrain Inflammation and Fibrosis. Cell Mol Gastroenterol Hepatol 2023;15:765–795.

38. Rueda GH, Causada-Calo N, Borojevic R, et al. Oral tryptophan activates duodenal aryl hydrocarbon receptor in healthy subjects: a crossover randomized controlled trial. Am J Physiol Gastrointest Liver Physiol 2024;326:G687–G696.

39. Huang Z, de Vries S, Fogliano V, et al. Effect of whole foods on the microbial production of tryptophan-derived aryl hydrocarbon receptor agonists in growing pigs. Food Chem 2023;416:135804.

40. Haq S, Wang H, Grondin J, et al. Disruption of autophagy by increased 5-HT alters gut microbiota and enhances susceptibility to experimental colitis and Crohn’s disease. Sci Adv 2021;7:eabi6442.

41. Huang Z, Boekhorst J, Fogliano V, et al. Impact of High-Fiber or High-Protein Diet on the Capacity of Human Gut Microbiota To Produce Tryptophan Catabolites. J Agric Food Chem 2023;71:6956–6966.

42. Rondeau LE, Da Luz BB, Santiago A, et al. Proteolytic bacteria expansion during colitis amplifies inflammation through cleavage of the external domain of PAR2. Gut Microbes 2024;16:2387857.

43. Galipeau HJ, Caminero A, Turpin W, et al. Novel Fecal Biomarkers That Precede Clinical Diagnosis of Ulcerative Colitis. Gastroenterology 2021;160:1532–1545.

44. Natividad JM, Agus A, Planchais J, et al. Impaired Aryl Hydrocarbon Receptor Ligand Production by the Gut Microbiota Is a Key Factor in Metabolic Syndrome. Cell Metab 2018;28:737–749 e4.

45. Hayes CL, Dong J, Galipeau HJ, et al. Commensal microbiota induces colonic barrier structure and functions that contribute to homeostasis. Sci Rep 2018;8:14184.

46. Santiago A, Hann A, Constante M, et al. Crohn’s disease proteolytic microbiota enhances inflammation through PAR2 pathway in gnotobiotic mice. Gut Microbes 2023;15:2205425.

47. Galipeau HJ, Rulli NE, Jury J, et al. Sensitization to gliadin induces moderate enteropathy and insulitis in nonobese diabetic-DQ8 mice. J Immunol 2011;187:4338–46.

48. Kechin A, Boyarskikh U, Kel A, et al. cutPrimers: A New Tool for Accurate Cutting of Primers from Reads of Targeted Next Generation Sequencing. J Comput Biol 2017;24:1138–1143.

49. Callahan BJ, McMurdie PJ, Rosen MJ, et al. DADA2: High-resolution sample inference from Illumina amplicon data. Nat Methods 2016;13:581–3.

50. Price MN, Dehal PS, Arkin AP. FastTree 2--approximately maximum-likelihood trees for large alignments. PLoS One 2010;5:e9490.

51. McMurdie PJ, Holmes S. phyloseq: an R package for reproducible interactive analysis and graphics of microbiome census data. PLoS One 2013;8:e61217.

