## Supplement for "Dietary tryptophan enhances aryl hydrocarbon receptor activation and reduces colitis through microbial metabolism"

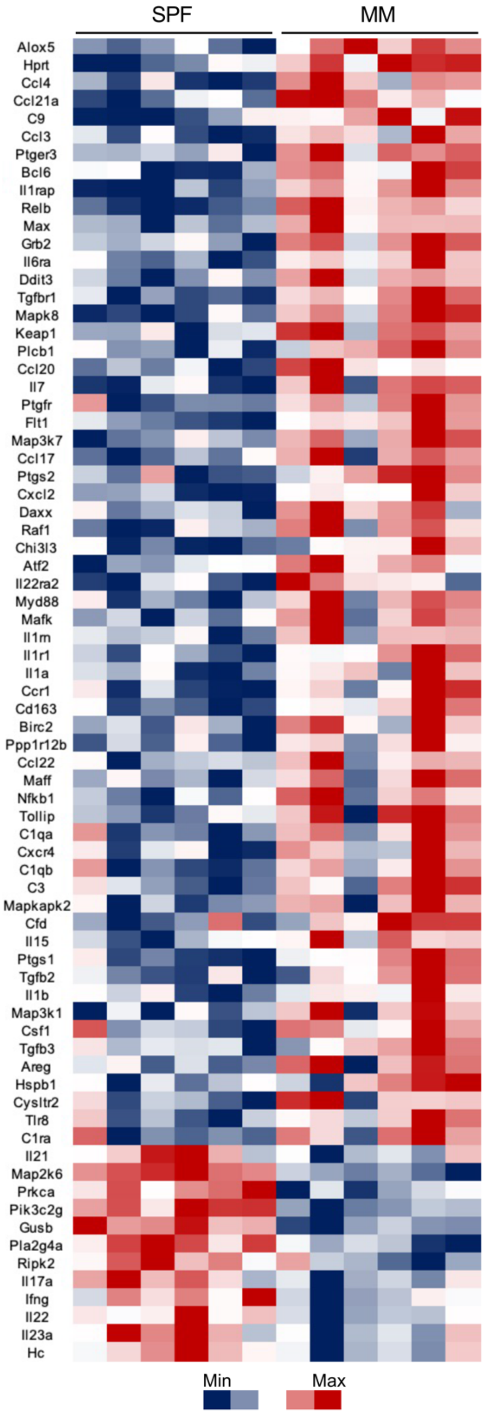


**Figure S1. Reduced microbial tryptophan metabolism increases pro-inflammatory gene expression during colitis**. Heat map of gene expression in the colon of specific pathogen-free (SPF) and minimal microbiota (MM)-colonized mice following DSS-induced colitis. Only genes significantly dysregulated between groups are shown. MM-colonized mice, which exhibit reduced microbial tryptophan metabolism compared to SPF mice, show upregulation of pro-inflammatory genes (e.g., *Il1b*, *Tnf*) and downregulation of regulatory or protective genes such as *Il22*. Colours represent normalized expression values: red indicates upregulation, and blue indicates downregulation. P-values were calculated using nSolver 2.5 software with the Wald test. (n=6 per group).

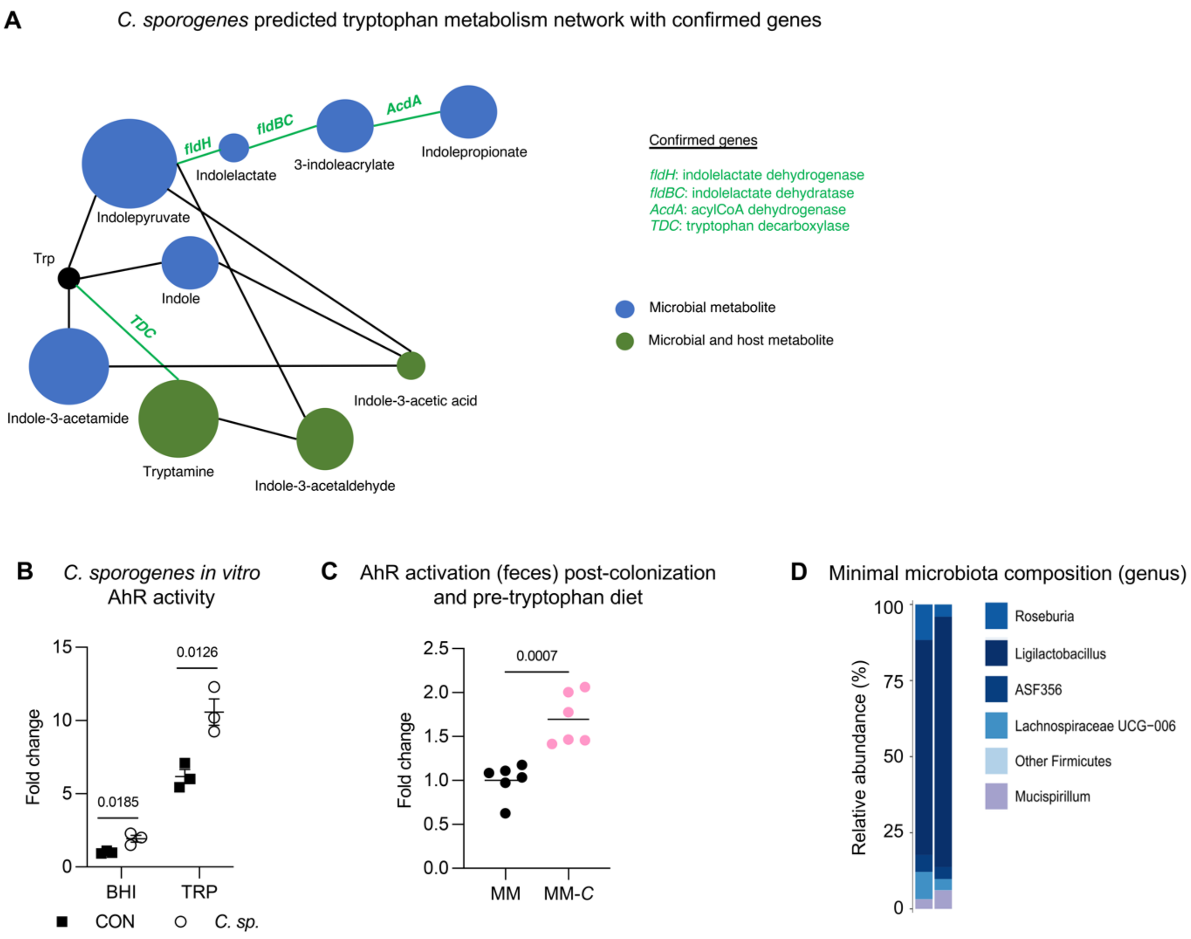


**Figure S2. *Clostridium sporogenes* isolate metabolizes tryptophan and increases AhR activation in mice. (a)** Predicted tryptophan metabolism network of *C. sporogenes*, with confirmed genes identified through whole genome sequencing highlighted in green. Metabolites include microbial-specific products (blue) and shared microbial and host metabolites (green). Green lines indicate confirmed pathways in *C. sporogenes*, while black lines indicate pathways predicted but not confirmed in this strain. Circle size reflects the predicted enzymatic activity for each metabolite. Confirmed genes include *fldH* (indolelactate dehydrogenase), *fldBC* (indolelactate dehydratase), *AcdA* (acyl-CoA dehydrogenase) and TCD (tryptophan decarboxylase; pyridoxal 5'-phosphate-dependent carboxylase). **(b)** AhR activation by *C. sporogenes* culture supernatants *in vitro*. Supernatants were collected from *C. sporogenes* grown in brain heart infusion (BHI) or tryptophan-supplemented (TRP) medium. (n=3) **(c)** AhR activation (feces) of minimal microbiota (MM) and minimal microbiota + *C. sporogenes* (MM-*C*) colonized mice after colonization and before tryptophan diet intervention. Data are presented as mean where each dot represent one mouse (n=6 per group). Statistical analysis was performed using an unpaired *t* test. **(d)** Genus level composition of minimal microbiota mouse intestinal content (n=2 mice).

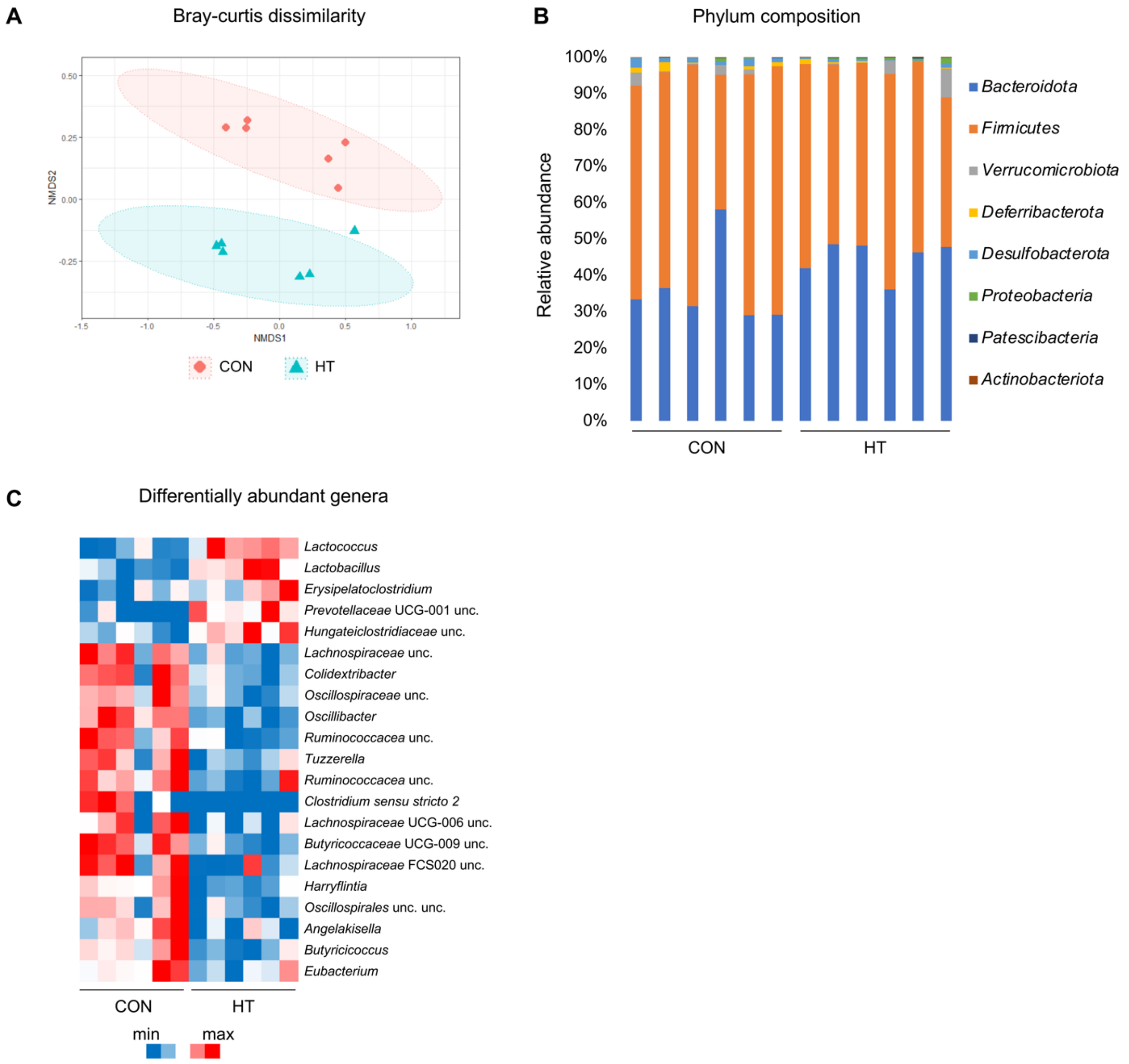


**Figure S3. Microbiota alterations in specific-pathogen-free mice after high tryptophan diet. (a)** Bray-Curtis dissimilarity showing distinct clustering of colonic microbiota of specific mice fed a control (CON) or high tryptophan diet (HT). **(b)** Relative abundance of bacterial phyla in colon content of CON- and HT-fed mice. **(c)** Heatmap of differentially abundant genera between CON and HT groups. Scale bar represents min to max values of relative abundance per genus. Statistical analyses for differentially abundant genera ($p < 0.05$) were performed using Mann-Whitney U tests.

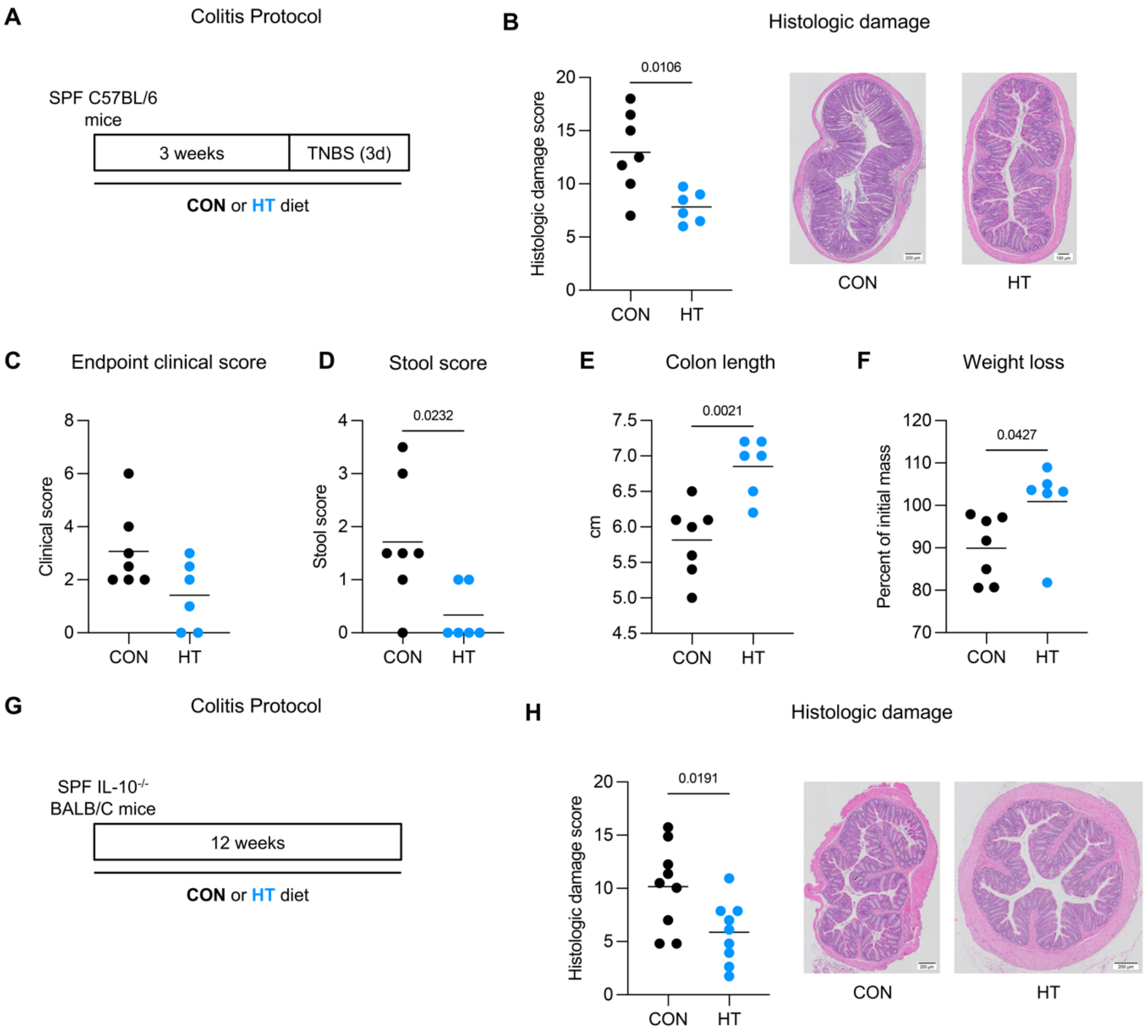


**Figure S4. Dietary tryptophan reduces hapten-induced and genetic colitis severity**. **(a)** Colitis protocol: Specific pathogen-free (SPF) C57BL/6 mice were fed a control (CON) or high tryptophan (HT) diet for three weeks, followed by induction of colitis using TNBS (2%) for three days. **(b)** Histologic damage scores of colon tissue with representative images (H&E-stained; scale bars = 100 μm). **(c)** Endpoint clinical scores. **(d)** Stool scores. **(e)** Colon length at endpoint. **(f)** Weight loss as a percentage of mass at initiation of TNBS colitis. **(g)** Colitis protocol: SPF IL-10$^{-}$/$^{-}$ BALB/c mice were fed a CON or HT diet for 12 weeks to assess spontaneous colitis severity. **(h)** Histologic damage scores of colon tissue with representative images (H&E-stained; scale bars = 200 μm). Data are presented as mean where each dot represents one mouse. Statistical analyses were performed using unpaired *t* tests.

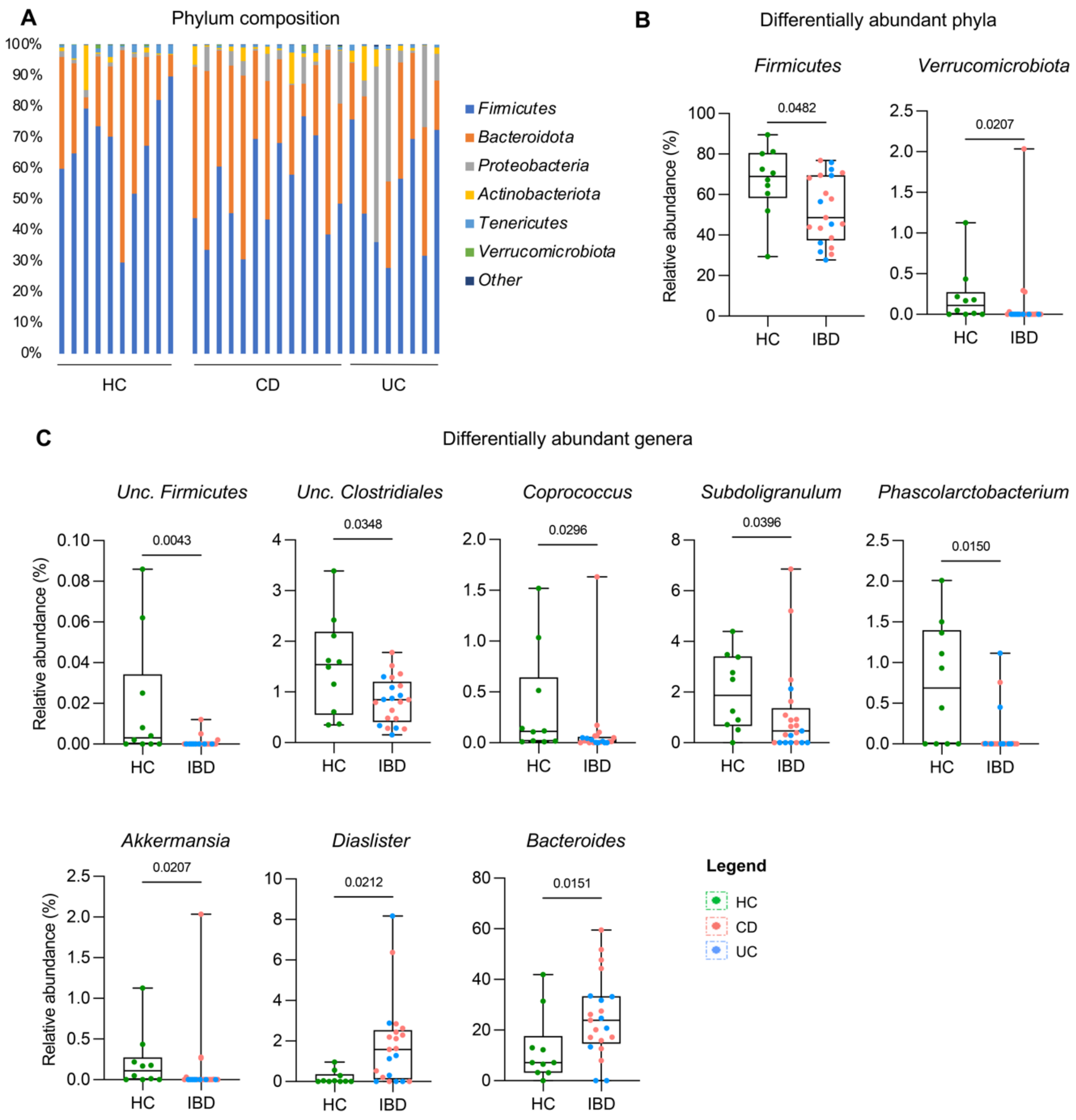


**Figure S5. IBD patients have altered microbiota compared to healthy controls. (a)** Phylum composition of fecal gut microbiota in healthy controls (HC), Crohn's disease (CD), and ulcerative colitis (UC) patients. Each bar represents one human sample. **(b)** Relative abundance of differentially abundant phyla comparing HC and inflammatory bowel disease patients (IBD). **(c)** Relative abundance of genera comparing HC and IBD patients. Data are presented as median with interquartile range with whiskers extending to the min and max data points, and where each dot represents one sample (HC, green; CD, pink; UC, blue). Statistical analysis was performed with Mann-Whitney U tests.

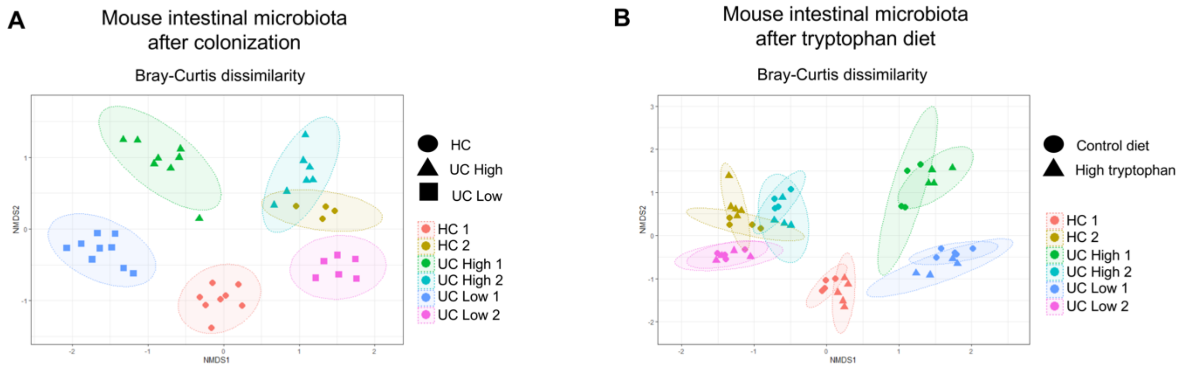


**Figure S6. Microbiota after transfer from IBD patients and healthy controls to germ-free mice and after tryptophan supplementation. (a)** Bray-Curtis dissimilarity of mouse intestinal microbiota after colonization, showing clustering based on donor microbiota (HC, UC High, or UC Low). **(b)** Bray-Curtis dissimilarity of mouse intestinal microbiota after colonization and CON or HT diet, showing clustering based on donor microbiota and diet conditions.

**Table S1.** Patient and healthy control demographic information.

| Status | Diagnosis | Sex | Age | Race | Past medical history | Medication |
|---|---|---|---|---|---|---|
| IBD | CD | M | 65.0 | Caucasian | cerebellar ataxia, fistulotomy (59 y/o) | 5-ASA (Salofalk 1 g/d); rabeprazole 20 mg/d; Vit B12 |
| | CD | F | 68.0 | Caucasian | ileostomy and bowel resection (56 y/o) | 5-ASA |
| | CD | F | 66.0 | Caucasian | cardiovascular disease; allergies; C-section | Asacol 800 mg/day; omeprazole 20 mg/d; adalat PRN |
| | CD | M | 53.0 | Caucasian | none | none |
| | CD | M | 35.0 | Caucasian | bowel resection (13 y/o) | pentasa |
| | CD | F | 32.0 | Caucasian | none | none |
| | CD | F | 48.0 | Caucasian | none | salofalk |
| | CD | M | 28.0 | Caucasian | none | none |
| | CD | F | 53.0 | Caucasian | none | salofalk |
| | UC | F | 47.0 | Caucasian | none | asacol |
| | UC | F | 60.0 | Caucasian | cutaneous sarcoidosis, hypertension, cholecystectomy | salofalk, buscopan |
| | UC | F | 49.0 | Caucasian | psoriasis, osteopenia, graves disease, celiac disease | none |
| | UC | F | 67.0 | Caucasian | hypertension, hiatial hernia repair surgery, polectomy | salofalk, asacol, aspirin, hydrochlorothiazide |
| | UC | M | 52.0 | Caucasian | none | asacol |
| | UC | F | 51.0 | Caucasian | headaches, obstructive sleep apnea | none |
| | UC | F | 53.0 | Caucasian | appendectomy in 2012 | asacol |
| | UC | F | 23.0 | Caucasian | none | prednisone |
| | UC | M | 22.0 | Caucasian | Primary sclerosing cholangitis | ursodiol, mezavant |
| | UC | M | 71.0 | Caucasian | none | 5-ASA |
| | UC | F | 64.0 | Caucasian | none | asacol, immuran |
| | UC | F | 66.0 | Caucasian | hypertension | coversyl 40 mg daily, meloxicam |
| | UC | F | 64.0 | Caucasian | glaucoma, eczema | glaucoma drops |
| Healthy | Healthy | F | 39.0 | Caucasian | GERD, colonic and gastric polyps, arthritis | pantoprazole 40 mg daily |
| | Healthy | F | 61.0 | Caucasian | endometriosis, tinnitus | omega 3, calcium |
| | Healthy | M | 60.0 | Caucasian | iron deficiency anemia | none |
| | Healthy | F | 59.0 | Caucasian | hypertension | coversyl 40 mg daily |
| | Healthy | F | 53.0 | Caucasian | hypertension | hydrochlorothiazide |
| | Healthy | M | 63.0 | Caucasian | hypertension | diovan 160 mg OD |
| | Healthy | F | 45.0 | Indian | hypertension during pregnancy; dyslipidemia, anemia, thyroidectomy, c-section | levothyroxine, iron, multivitamins |
| | Healthy | M | 31.0 | Caucasian | GERD, asthma | ventolin |
| | Healthy | F | 69.0 | Caucasian | hypertension, hemorrhoids, Hypothyroidism | synthroid, atenolol, meloxicam |
| | Healthy | F | 34.0 | Caucasian | anemia, migraines, celiac disease | iron |

**Table S2.** Tryptophan diet composition.

| | Control diet TD.00102 | High tryptophan diet TD.210165 |
|---|---|---|
| **Nutrient, % by weight** | | |
| Protein | 12.4 | 13.2 |
| Fat | 4.1 | 4.3 |
| Carbohydrate | 68.4 | 67.6 |
| **Nutrient, % kcal from** | | |
| Protein | 13.8 | 14.6 |
| Fat | 10.3 | 10.3 |
| Carbohydrate | 75.9 | 75.1 |
| kcal/g | 3.6 | 3.6 |
| **Ingredients, g/kg** | | |
| Casein | 140 | 140 |
| L-Cystine | 1.8 | 1.8 |
| L-Tryptophan | 0 | 8 |
| Corn Starch | 460.942 | 452.892 |
| Maltodextrin | 100 | 135 |
| Sucrose | 150 | 150 |
| Soybean Oil | 40 | 40 |
| Cellulose | 50 | 50 |
| Vitamin Mix, AIN-93-VX | 14.5 | 14.5 |
| Choline Bitartrate | 2.75 | 2.75 |
| TBHQ, antioxidant | 0.008 | 0.008 |
| Mineral Mix, AIN-93M-MX (94049) | 35 | 35 |
| Food colouring | | 0.05 |
| **Mineral Composition, g/kg** | | |
| Calcium | 5 | 5 |
| Phosphorus | 3 | 3 |
| Potassium | 3.6 | 3.6 |
| Sodium | 1 | 1 |
| Chlorine | 1.6 | 1.6 |
| Magnesium | 0.517 | 0.517 |
| Copper | 6.1 | 6.1 |
| Iron | 36.9 | 36.9 |
| Zinc | 39.8 | 39.8 |
| Manganese | 10.5 | 10.5 |
| Iodine | 0.21 | 0.21 |
| Selenium | 0.15 | 0.15 |
| Molybdenum | 0.15 | 0.15 |
| Chromium | 1 | 1 |

**Table S3.** Primer sequences for RT-qPCR.

| | | |
|---|---|---|
| *Ahr* | Forward | 5'-GAGCTTCTTTGATGGCGCTG-3' |
| | Reverse | 5'-GTCCACTCCTTGTGCAGAGT-3' |
| *Ahrr* | Forward | 5'-CCATTCAGAAGCGCCTTGCAG-3' |
| | Reverse | 5'-AGGCAGCGAACACGACAAAT-3' |
| *Cyp1a1* | Forward | 5'-ACATTGTGCCTGCCTCCTAC-3' |
| | Reverse | 5'-GTAGGGTGAACAGAGGTGCC-3' |
| *Il22* | Forward | 5'-CATGCAGGAGGTGGTACCTT-3' |
| | Reverse | 5'-CAGACGCAAGCATTTCTCAG-3' |
| *Gapdh* | Forward | 5'-AACTTTGGCATTGTGGAAGG-3' |
| | Reverse | 5'-ACACATTGGGGGTAGGAACA-3' |
